# An artificial metal-free peroxidase designed using a ferritin cage

**DOI:** 10.1101/2025.02.11.637337

**Authors:** Jiaxin Tian, Basudev Maity, Tadaomi Furuta, Tiezheng Pan, Takafumi Ueno

## Abstract

Developing artificial enzymes is challenging because it requires precise design of active sites with well-arranged amino acid residues. Histidine-rich oligopeptides have been recently shown to exhibit peroxidase-mimetic activities, but their catalytic function relies on maintaining unique supramolecular structures. This work demonstrates the design of a specific array of histidine residues on the internal surface of the ferritin cage to function as an active center for catalysis. The crystal structures of the ferritin mutants revealed histidine-histidine interactions, forming well-defined histidine clusters (His-clusters). These mutants exhibit peroxidase-mimetic activities by oxidizing 3,3’,5, 5’-tetramethylbenzidine (TMB) in the presence of hydrogen peroxide. Molecular dynamics simulations further highlight the co-localization of TMB and hydrogen peroxide at the histidine-rich clusters, indicating that the confined environment of the ferritin cage enhances their interactions. This study presents a simple yet effective approach to design cofactor-free artificial enzymes, paving the way for innovations in bioinspired catalysis.

## Introduction

The catalytic function of enzymes arises from active sites formed through the cooperation of cofactors and functional groups. To mimic the active sites of natural enzymes, a common approach involves introducing key functional groups and cofactors to create active pockets.^1,2^ Self-assembled peptides without typical cofactors have been recently shown to exhibit various enzyme-mimetic activities, including oxidase, reductase, and esterase activities.^3, 4, 5, 6^ Specific arrangements of functional residues, such as histidine or the formation of the catalytic dyad,^7^ and triad,^8^ drive these activities. While such cofactor-free nanocatalysts are promising, their reliance on higher-order nanostructures limits the flexibility in designing enzyme-mimetic catalysts. Thus, there is significant interest in systematically introducing functional residues into self-assembled proteins. Based on this rationale, we selected protein cages^9^ as suitable templates, given their ability to self-assemble from protein monomers. In this study, we focused on introducing amino acid clusters onto the internal surface of protein cages to promote catalytic reactions.

Self-assembled protein cages form highly organized and symmetric nanostructures with large internal cavities.^9, 10^ These cavities create unique microenvironments that can isolate molecules and provide confined spaces for chemical reactions.^11^ This contrasts with catalytic peptide nanostructures, where only the outer surfaces with suitable arrangements of functional residues are available for reactions.^12^ However, most research on protein cages has focused on immobilizing foreign enzymes,^13,14^ and cofactors^15^ for catalytic reactions, rather than repositioning specific residues to create new active centers. Therefore, the primary goal of this study is to align functional amino acids, such as histidine, on the internal surface of protein cages to form clusters that can exhibit enzyme-mimetic activities while leveraging the structural advantages of the protein cage. Achieving optimal residue alignment through systematic design is crucial for enhancing catalytic activities. For this purpose, we selected the ferritin cage as our model system.

Among various protein cages, ferritin, whose 24 subunits form a hollow cage with inner and outer diameters of 8 and 12 nm, respectively, is quite robust and valuable for biomaterial applications.^16^ The biological role of ferritin is to function as an iron storage protein. The mechanism by which ferritin accumulates iron involves the oxidation of incorporated Fe(II) to Fe(III) by hydrogen peroxide, synthesized at the oxidase center within the cage, which oxidizes incorporated Fe(II) to Fe(III).^17^ The oxidase centers include specific amino acids located near the two-fold symmetric interface.^18^ By substituting suitable amino acids such as cysteine and histidine, various metal binding sites, including organometallic Ir,^19^, ^15^Rh,^20^ and Ru complexes,^21^ exhibit catalytic and biological functions in various reactions. However, apart from such properties of metal-active centers, engineered catalytic reactions promoted by specific placement of specific amino acid residues have not been achieved. Recently, we reported the construction of aromatic clusters at the two-fold symmetric interfaces by introducing multiple phenylalanine (Phe) residues.^22^ This modification was found to enhance the thermal stability of the cage without affecting the 24-mer cage formation. Based on these findings, it is expected that histidine (His)-clusters can be constructed by systematically introducing multiple His residues on the B-helix at the two-fold symmetric interface (Figure 1a).^23,24^ While His-clusters are recognized for their roles in metal binding and other biological functions,^23,25^ the objective of the present work was to develop these clusters as catalytic centers within the confined environment of the ferritin cage.

**Figure 1:**
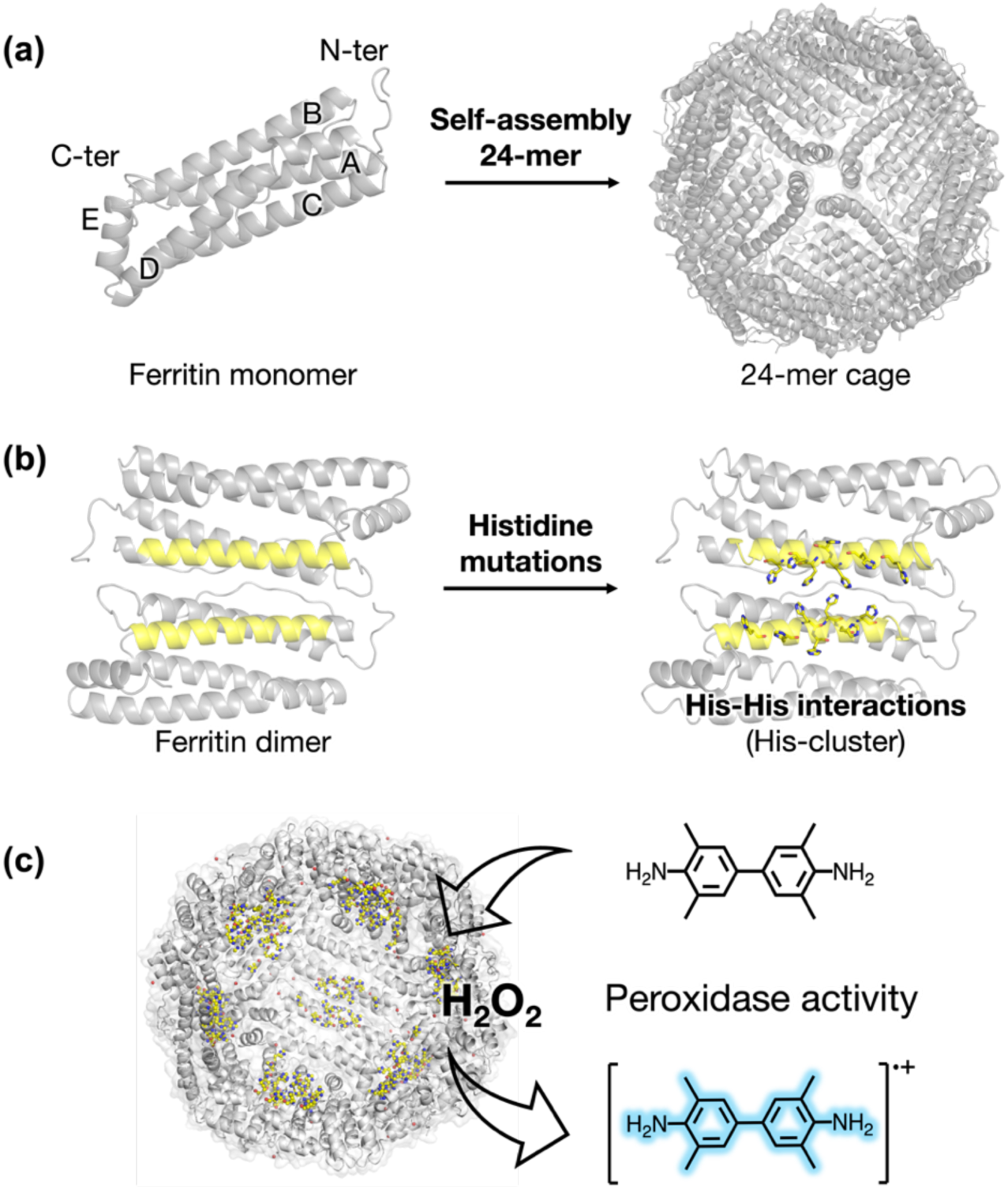
Design of His-cluster in ferritin inspired by cofactor/metal free self-assembly peptide. (a) Structure of FrWT cage formed by the self-assembly from single subunit consisting of 4 helices (A-D) and a short E helix. (b) The His-cluster design position at the two-fold symmetric interface. (c) Working model of construction of His-cluster in ferritin and the corresponding peroxidase mimetic activity in the presence of H_2_O_2_.

Herein, we report the construction of a His-cluster aligned along the two-fold symmetric interface of the ferritin cage and evaluate the peroxidase mimetic activities promoted by the clusters (Figure 1). We confirmed the formation of the His-clusters in high-resolution crystal structures. The artificially constructed His-cluster in the ferritin cage with six mutated His residues promotes very high catalytic one-electron oxidation of 3, 3’, 5, 5’-tetramethylbenzidine (TMB) substrate in the presence of hydrogen peroxide (H_2_O_2_). The catalytic efficiency is over 80-fold higher than that of an oligohistidine nanostructure containing 15 His residues (Table 2).^26^ By controlling the positions and number of His residues within the His-cluster, we investigated the role of His-cluster as a catalytic center in the ferritin cage and characterized its structure-activity relationship. Molecular dynamic (MD) simulations revealed simultaneous interactions of TMB and H_2_O_2_ on the His-cluster which are influenced by the effect of the confined protein cage environment. The molecular design presented here effectively utilizes the structural integrity of ferritin, positioning active sites at the two-fold symmetric interface without compromising the overall architecture, thereby indicating new possibilities for using protein cages in catalysis.

## Results

### Design and preparation of histidine clusters in a ferritin cage

We chose recombinant wild-type L-chain ferritin from horse spleen (FrWT) as a suitable scaffold to design His-clusters because of its unique structural features, especially the symmetric interfaces (Figure 1c). The two-fold symmetric interface is known to be the most stable section due to its large contact area compared to other subunit interfaces.^27,28^ Previously, we introduced multiple Phe residues on the B-helix at the two-fold symmetric axis to construct aromatic clusters, improving their thermal stability with retention of the 24-mer ferritin cage.^22^ Aromatic clusters with benzene ring (Phe) distances of approximately 5-6 Å were formed between Phe residues on adjacent helices. Similarly, amino acid mutations can be made on adjacent helices to create active centers. Since proximal His residues with hydrogen bond distances of about 3Å can activate H_2_O_2_ in peptide-based nanomaterials,^29^ our motivation is to construct His-clusters at the two-fold symmetric interface to develop artificial oxidase-mimetic catalysis (Figure 1b,c).

To construct the His-cluster at the two-fold symmetric interface, we primarily considered the C^α^-C^α^ distances between amino acids near R59, similar to the previous construction of Phe-clusters.^22^ In FrWT, from E53 to E63, the C^α^-C^α^ distances of two amino acids on adjacent helices range from 5.0 to 6.4 Å. When the residues E53, E57, R60, and R64 were mutated to Phe, the C^α^-C^α^ distances were found to remain similar (4.9 to 6.1 Å).^22^ Considering the smaller 5-membered imidazole ring (His) and the H-bond donor/acceptor properties, a His-cluster is expected to form at those positions if histidines are introduced.^23,24^ Therefore, we designed a series of mutants containing His with varying numbers from 2 to 8. These mutants are designated **Fr-H2**(E53H/E57H), **Fr-H4ex**(E53H/E57H/E60H/R64H), **Fr-H4in**(R52H/E56H/R59H/E63H), **Fr-H6**(E53H/E56H/E57H/R59H/E60H/E63H) and **Fr-H8**(R52H/E53H/E56H/E57H/R59H/E60H/E63H/R64H (Table 1). The mutants were expressed in *E. coli* and purified by anion exchange and size-exclusion chromatography (See Supporting Information). Subsequently, the mutants were characterized by various analytical methods, including X-ray crystallography. The formation of the 24-mer cage was confirmed by size-exclusion chromatography (SEC), and native polyacrylamide gel electrophoresis (PAGE), which showed a band pattern similar to that of FrWT (Figure S1). The monomer masses of the mutants were confirmed by matrix-assisted laser desorption ionization-time of flight mass spectrometry (MALDI-TOF-MS) (Figure S1). The circular dichroism (CD) measurement followed by Native PAGE analysis indicates that the 24-mer cage structures of Fr-His remain as stable as FrWT (Figure S2).

**Table 1:** List of designed ferritin mutants and their sequences.^a^.

| Mutants | Residue positions from 48 to 64 |  |  |  |  |  |  |  |  |  |  |  |  |  |  |  |  |
| --- | --- | --- | --- | --- | --- | --- | --- | --- | --- | --- | --- | --- | --- | --- | --- | --- | --- |
|  | 48 | 49 | 50 | 51 | 52 | 53 | 54 | 55 | 56 | 57 | 58 | 59 | 60 | 61 | 62 | 63 | 64 |
| FrWT | C | H | F | F | R | E | L | A | E | E | K | R | E | G | A | E | R |
| Fr-H2 | C | H | F | F | R | **H** | L | A | E | **H** | K | R | E | G | A | E | R |
| Fr-H4in | C | H | F | F | **H** | E | L | A | **H** | E | K | **H** | E | G | A | **H** | R |
| Fr-H4ex | C | H | F | F | R | **H** | L | A | E | **H** | K | R | **H** | G | A | E | **H** |
| Fr-H6 | C | H | F | F | R | **H** | L | A | **H** | **H** | K | **H** | **H** | G | A | **H** | R |
| Fr-H8 | C | H | F | F | **H** | **H** | L | A | **H** | **H** | K | **H** | **H** | G | A | **H** | **H** |

<sup>a</sup>The mutated residues have been highlighted in bold.

### Crystal structures of ferritin mutants

To determine the atomic-level structures, we crystallized the ferritin mutants with varying numbers of His residues using a previously reported procedure and obtained X-ray diffraction measurements.^15^ ^30^ The high-resolution (1.5-1.6 Å) structures were determined using wild-type ferritin (PDB 1DAT) as a starting model. The crystallographic data and refinement statistics are listed in Table S2. All structures maintain the 24-mer cage structure with a root mean square deviation (C^⍺^ atoms) up to 0.17 Å compared to that of FrWT. The mutated His residues were observed clearly (Figure S3) with relatively high B-factors compared to surrounding residues (Figure S4. This implies that the His residues are flexible. We analyzed the His geometries at the two-fold symmetric interface and the His-His distances, which fall into the category of His-cluster as defined by H. Sun et al. (Table S3).^23^ The His residues participate in both inter-subunit and intra-subunit interactions (Figure S5). The His-His distances between 4.0 Å and 7.5 Å are particularly important because such pairs have a high probability of interacting with each other (Figure 2).^31^ In addition to the observation of His-His interactions, several water molecules were found to form hydrogen-bonding interactions with both intermolecular and intramolecular His residues (Figure S6).

**Figure 2.**
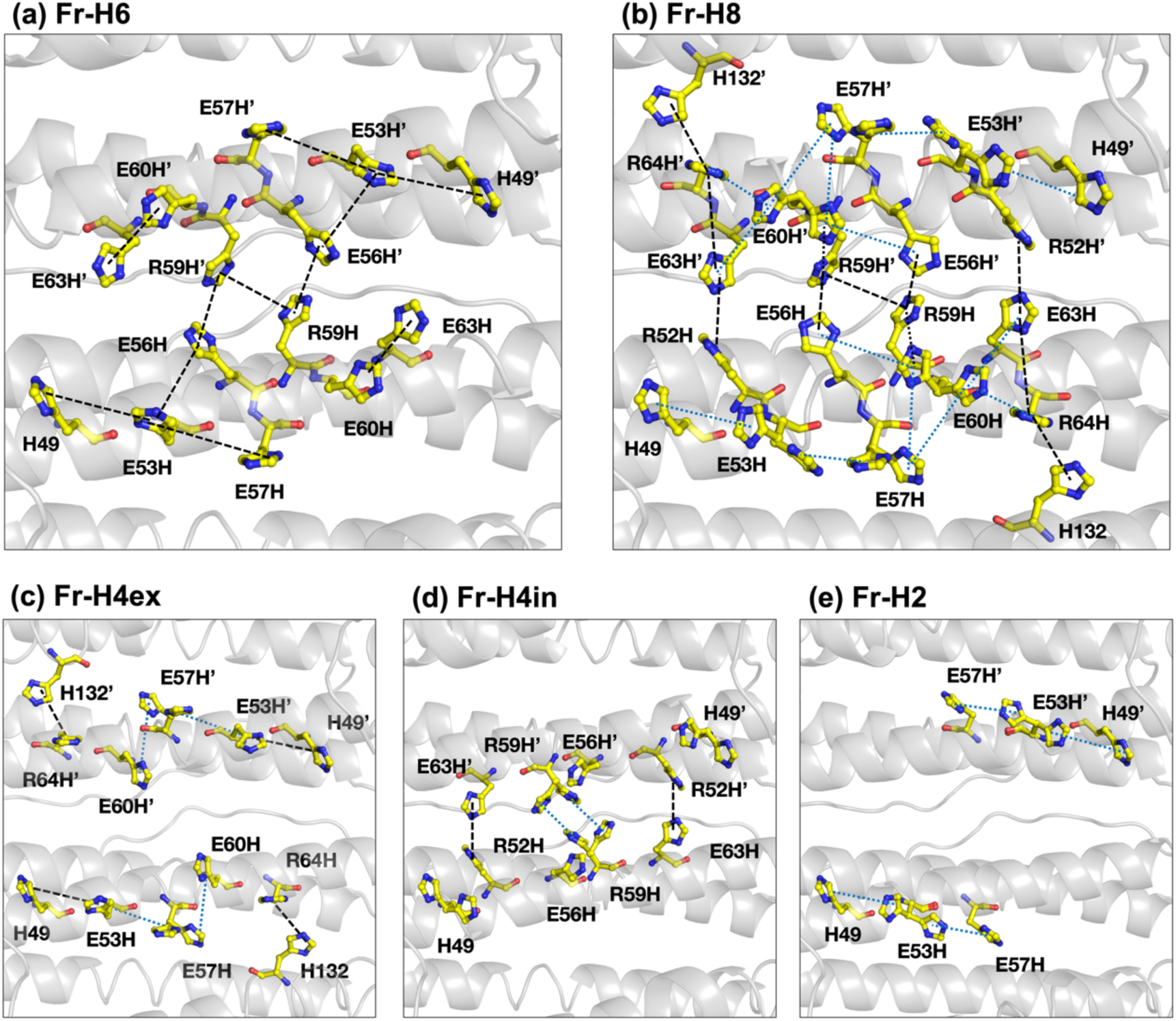
Crystal structure of ferritin His mutant. (a-e) Closeup view of the His site of **Fr-H8**, **Fr-H6**, **Fr-H4in**, **Fr-H4ex** and **Fr-H2**, respectively. The black dashed lines represent the interactions between the centers of the imidazole rings (centers) between two histidine. The cyan dotted lines show those interactions between the centers of the imidazole rings from Histidine with double conformation. His-His distances longer than 4 Å shorter than 7.5 Å are displayed.

In **Fr-H6**, the residues His49, His53 and His57 interact with each other in a row with ring-to-ring distances of 6.7 Å and 7.4 Å, respectably (Figure 2a and Table S3). Similarly, His60 and His63 also interact with each other with a ring-to-ring distance of 5.2 Å. The residues His56 and His59 are directed towards the two-fold interface, and thus, interact with each other as well as the corresponding His59’ and His56’ residues of the other subunit to participate in inter-subunit interactions. This long histidine network is formed starting with His49 of one monomer and extends to His49’ of the adjacent monomer. The ring-to-ring distances of His-His pairs range from 5.2 to 7.4 Å. In Fr-H8, two additional His residues are introduced at positions 52 and 64. This results in additional inter-subunit interactions involving these residues so that His52’, His63, His64 and His132 form a long network. In addition, in the Fr-H8 structure, binding interactions of His residues with several Cd ions (which are used as a crystallizing reagent) were observed, involving His53 and His57, His60 and His64 (Figure S6). Related to these observations, double conformations were observed for His53, His57 and His60, which were not observed in Fr-H6 structure. Such double conformations indicate the flexibility of these residues which provides additional interactions (Figure 2b). One conformer of His60 interacts with His63/64 and His57, and its other conformer interacts with His56/57 and His59. Similarly, two conformers of His57 interact separately with one conformer each of His60 and His53. The other His-His interactions remain the same as in Fr-H6. This suggests that Fr-H6 has a more compact and simpler arrangement His-His interactions than the more flexible and complicated arrangement of Fr-H8.

By reducing the number of His residues, the structures of **Fr-H4in**, **Fr-H4ex**, and **Fr-H2** showed distinct features. In all the cases, His-clusters are formed on the B-helix (Figure 2c-e). The mutants **Fr-H4in** and **Fr-H4ex,** which have the same number of His residues, showed intra-(**Fr-H4ex**) and inter-(**Fr-H4in**) subunit interactions among the His residues. In **Fr-H4in**, the His residues are located towards the inter-subunit interfaces causing His52, His59, and His63 to interact with His63’, His59’, and His52’ of the opposite subunit (Figure 2d) with His59 having a double conformation. In **Fr-H2**, the His-cluster consists of three His residues. His53 has a double conformation that generates interactions with His49 and His57 at distances of 6.2 Å and 5.6 Å, respectively (Figure 2e and Table S3).

### Peroxidase mimetic activities

We investigated the oxidase mimetic activities of the Fr-mutants using 3,3’,5,5’-tetramethylbenzidine (TMB) as a substrate, which is known to undergo one-electron oxidation catalyzed by peroxidase in the presence of H₂O₂.^26,29,32^ We first performed the reaction using **Fr-H8** which has the highest number of His residues. In the presence of H_2_O_2_, **Fr-H8** oxidizes TMB and generates the characteristic absorbance at 652 nm for the one-electron oxidation reaction which was not observed for FrWT (Figure 3a). This suggests that the His-cluster consisting of eight His residues in **Fr-H8** is responsible for the activity, while FrWT, which lacks a His-cluster, remains inactive under the same conditions. To avoid any possible metal contamination, all reactions were performed after purifying the mutants from a solution containing ethylenediaminetetraacetic acid (EDTA). The peroxidase activity of **Fr-H8** is highest at pH 4.0 and gradually decreases with increasing pH (Figure S7a). The same trend is followed for the **Fr-H6** mutant (Figure S7b).

**Figure 3.**
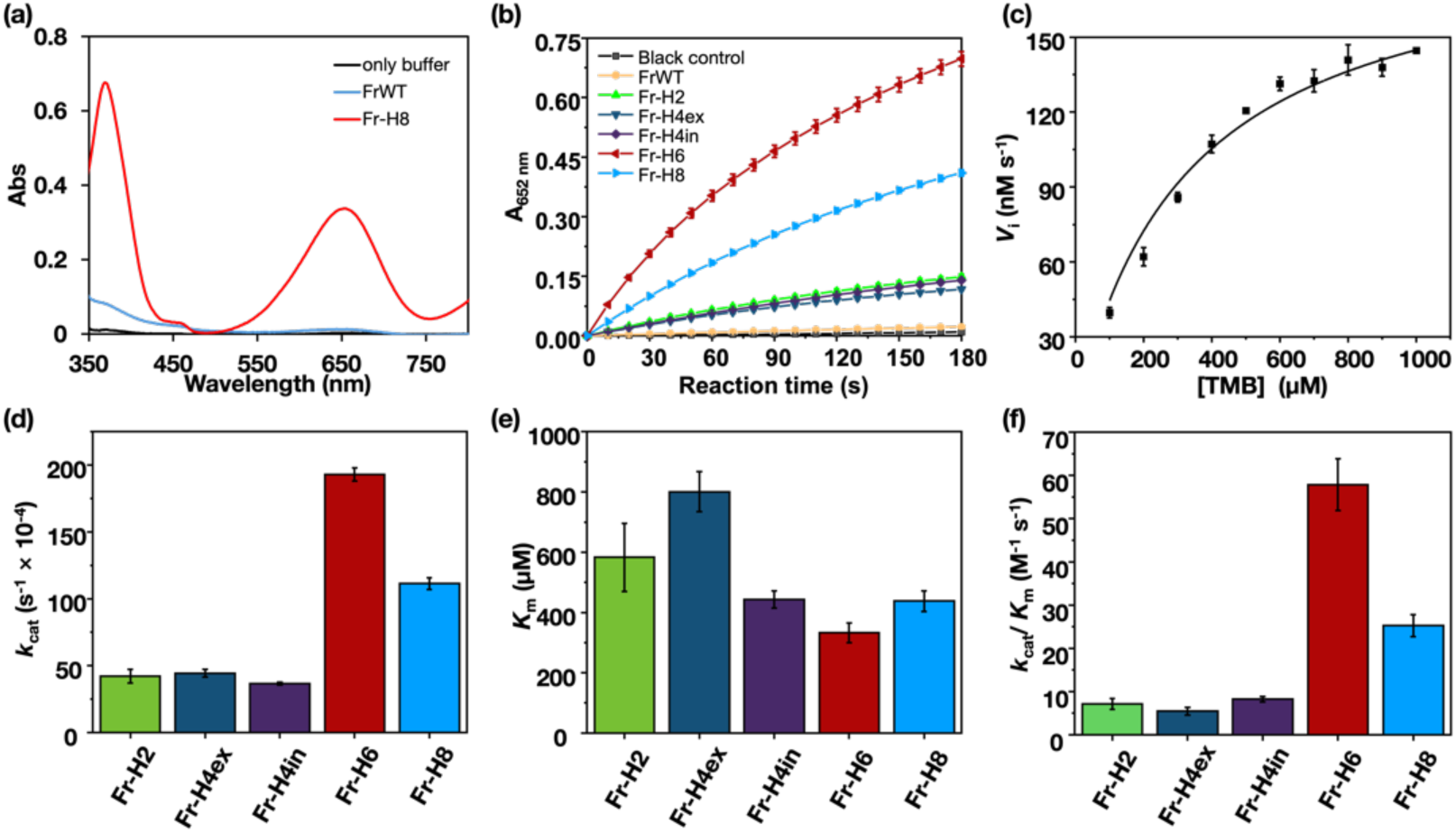
Evaluation of oxidase mimetic activities by Fr-His mutants. (a) Absorption spectrum of the TMB oxidation by **Fr-H8** in the presence of H_2_O_2_ compared to that of FrWT. Reaction condition: [protein] = 10 µM, [TMB] = 700 µM, [H_2_O_2_] = 20 mM in 50 mM NaOAc (pH 4.0). Reaction time: 3 min. (b) Time dependent absorbance changes at 652 nm for TMB oxidation FrWT and its mutants under the same conditions as in (a). (c) Changes of reaction velocity (*V*_i_) with increasing TMB concentration for the TMB oxidation by **Fr-H6**. [protein] = 10µM, [H_2_O_2_] = 20 mM. (d) *k*_cat_ (s^−1^×10^−4^), (e) *K*_m_ (µM) and (f) *k*_cat_/*K*_m_ (M^−1^s^−1^) for the Fr-His mutants determined from the Michaelis– Menten curve. [protein] = 10µM, [H_2_O_2_] = 20 mM.

We assessed the catalytic activities of the other mutants at pH 4.0. **Fr-H6** has the highest activity followed by **Fr-H8** (Figure 3d). **Fr-H4in**, **Fr-H4ex**, and **Fr-H2** have lower activity compared to **Fr-H6**. The activity increases as the number of mutated His residues increases but the relationship is not linear. This indicates that suitable positioning of His residues is essential for optimal activity (Figure 2, 3b and S5).

To investigate the reaction kinetics, we investigated the catalysis by varying the concentrations of TMB and H_2_O_2_ as substrates. The results follow typical Michaelis– Menten kinetics (Figure 3c-3f). The kinetic parameters *k*_cat_, *K*_m_, and *k*_cat_/*K*_m_ values are listed in Table 2. Among all mutants, **Fr-H6** has the lowest *K*_m_ value of 333.40 µM, indicating that this mutant has the strongest affinity for TMB. As a result, **Fr-H6** exhibits the most rapid turnover (*k*_cat_, 193.03×10^−4^ s^−1^) of TMB molecules per enzyme active site compared to other ferritin mutants. Among all mutants, **Fr-H6** has the highest catalytic efficiency (*k*_cat_/*K*_m_, 57.95) for TMB oxidation at about 82 times higher than that of oligohistidine (*k*_cat_/*K*_m_, 0.70).^26^ **Fr-H8** has about half of the catalytic efficiency (*k*_cat_/*K*_m_, 25.47) of **Fr-H6** and **Fr-H4in**, **Fr-H4ex**, and **Fr-H2** exhibit 7-10 times lower activity than **Fr-H6** but about ten times higher activity than that of oligohistidine (Table 2).^26^ The catalytic activities of the mutants follow the order of the *K*_m_ values, which are associated with substrate affinity for the catalysts.

**Table 2.** Comparison of kinetic parameters of TMB oxidation by Ferritin mutants.

| Catalysis | Substrate | $k_{\text{cat}}$ ($\text{s}^{-1}$) | $K_{\text{m}}$ (mM) | $k_{\text{cat}}/K_{\text{m}}$ ($\text{M}^{-1} \text{s}^{-1}$) |
| --- | --- | --- | --- | --- |
| FrWT | TMB | ND | ND | ND |
| **Fr-H2** | TMB | $(42.11 \pm 5.11) \times 10^{-4}$ | $0.58 \pm 0.11$ | $7.22 \pm 1.24$ |
| **Fr-H4ex** | TMB | $(44.30 \pm 2.95) \times 10^{-4}$ | $0.80 \pm 0.07$ | $5.53 \pm 0.89$ |
| **Fr-H4in** | TMB | $(36.58 \pm 1.12) \times 10^{-4}$ | $0.44 \pm 0.03$ | $8.25 \pm 0.61$ |
| **Fr-H6** | TMB | $(193.03 \pm 4.99) \times 10^{-4}$ | $0.33 \pm 0.03$ | $57.95 \pm 6.00$ |
| **Fr-H8** | TMB | $(111.50 \pm 4.43) \times 10^{-4}$ | $0.44 \pm 0.03$ | $25.47 \pm 2.57$ |
| oligohistidine<sup>26</sup> | TMB | $(1.32 \pm 0.05) \times 10^{-4}$ | $0.19 \pm 0.02$ | $0.70 \pm 0.07$ |
| FrWT | H<sub>2</sub>O<sub>2</sub> | ND | ND | ND |
| **Fr-H2** | H<sub>2</sub>O<sub>2</sub> | $(45.12 \pm 2.55) \times 10^{-4}$ | $15.46 \pm 1.38$ | $0.29 \pm 0.03$ |
| **Fr-H4ex** | H<sub>2</sub>O<sub>2</sub> | $(32.99 \pm 0.77) \times 10^{-4}$ | $11.48 \pm 0.53$ | $0.29 \pm 0.01$ |
| **Fr-H4in** | H<sub>2</sub>O<sub>2</sub> | $(35.64 \pm 2.62) \times 10^{-4}$ | $10.00 \pm 2.10$ | $0.36 \pm 0.08$ |
| **Fr-H6** | H<sub>2</sub>O<sub>2</sub> | $(163.78 \pm 3.11) \times 10^{-4}$ | $4.16 \pm 0.29$ | $3.93 \pm 0.28$ |
| **Fr-H8** | H<sub>2</sub>O<sub>2</sub> | $(92.47 \pm 1.61) \times 10^{-4}$ | $7.83 \pm 0.28$ | $1.18 \pm 0.05$ |
| oligohistidine<sup>26</sup> | H<sub>2</sub>O<sub>2</sub> | $(1.15 \pm 0.05) \times 10^{-4}$ | $3.67 \pm 0.54$ | $0.03 \pm 0.46$ |

Note. Reaction condition: For TMB substrate: [protein] = 10 $\mu\text{M}$ , [H<sub>2</sub>O<sub>2</sub>] = 20 mM, [TMB] = 0.1-1.0 mM; For H<sub>2</sub>O<sub>2</sub> substrate: [protein] = 10 $\mu\text{M}$ , [TMB] = 0.7 mM, [H<sub>2</sub>O<sub>2</sub>] = 2.5-20 mM. The data for oligohistidine was obtained from ref 26. ND: no detectable activity.

When the Michaelis-Menten kinetics analysis was performed using H_2_O_2_ as a substrate, the catalytic performance of the ferritin mutants was found to follow the same trend observed for TMB (Table 2). The *K*_m_ values when TMB as used as a substrate were significantly lower (∼13 times for **Fr-H6**) than when H_2_O_2_ was used as a substrate. This suggests that the His-clusters have a stronger affinity for TMB than for H_2_O_2_. As a result, higher catalytic turnover and efficiency was observed for all mutants in the reaction with TMB compared to the reaction with H_2_O_2_ as substrate.

Interestingly, **Fr-H2** which has only three His residues (His49, His53, His57), has 10-fold higher catalytic activity than the previously reported oligohistidine containing 15 His residues but is 14 times less active than **Fr-H6**.^26^ The activity of **Fr-H2** is similar to that of **Fr-H4in** and **Fr-H4ex** (Figure. 3a).

The mechanistic pathway was evaluated in a radical scavenging experiment.^33^ Terephthalic acid (TA) is known to scavenge hydroxyl radicals by forming 2-hydroxy terephthalic acid.^34^ If the reaction proceeds via the hydroxyl radical pathway, the reaction rate should decrease in the presence of TA. When the peroxidase activity of **Fr-H6** was examined in the presence of TA, a characteristic emission band at 425 nm was observed, suggesting the formation of 2-hydroxy TA (Figure S9a). Then, the absorbance at 652 nm (TMB) was monitored and found to decrease with increasing TA concentration (Figure S9b). This was also reflected in reaction velocity (*V*_i_), which gradually decreases (Figure S9c). These results primarily suggest the involvement of hydroxyl radicals in the catalytic reaction, an observation which is consistent with the results of a previous report indicating that hydroxyl radicals are produced from a His-based amyloid-like fibril nanostructure.^29^

When we tested the possibility of catalase activity being generated by **Fr-H6** upon mixing with H_2_O_2_, bubble formation was observed.^29^ This observation suggests that H_2_O_2_ is decomposing in presence of **Fr-H6**, leading to the production of oxygen gas (Figure S10a). To analyze this further, we added TMB to the mixture at various intervals after addition of H_2_O_2_ to **Fr-H6**, and monitored the TMB oxidation activity (Figure S10b). The gradual decrease of the oxidized TMB product with absorbance at 652 nm suggests that the H_2_O_2_ concentration decreases over time due to the presence of catalase activity (Figure S10b), which is consistent with a previous report.^26^

### Dynamics of hydrogen peroxide inside the ferritin cage

To explore the dynamics of H_2_O_2_ in the ferritin cage, we performed MD simulations and compared the behavior of H_2_O_2_ in the 24-mer cage and the half-open cage (12-mer, quasi-planar) of **Fr-H6** (Figure 4a). The radial distribution function (RDF) of H_2_O_2_ from the H59 pair in the full cage is about 3-fold higher than that in the half-open cage. In other words, compared to the planar structure, when H_2_O_2_ is confined in a spherical cage, it has a high probability of accessing the His-cluster as a catalytic site, leading to high activity. This suggests that the high activity is due to the cage effect. Moreover, it was observed that H_2_O_2_ frequently contacts the His-clusters with repeated binding and dissociation through the simulation, occasionally being located between His pairs (Movie S1 and Figure S11). Focusing on the three dimers (AB, CD, EF), H_2_O_2_ molecules were identified between His63^A^ and His60^A^ at 61.4 ns, between His57^C^ and His53^C^ at 75.9 ns, and between His60^C^ and His56^C^ at 82.5 ns (Figure 4b and Movie S2). On the open surface of oligohistidine, it is expected that once H_2_O_2_ dissociates from a His pair, it will take longer to make contact again. On the other hand, in an isolated environment such as ferritin cage, it appears that the internal H_2_O_2_ can frequently contact the His-clusters as a result of the constricted cage effect.

**Figure 4:**
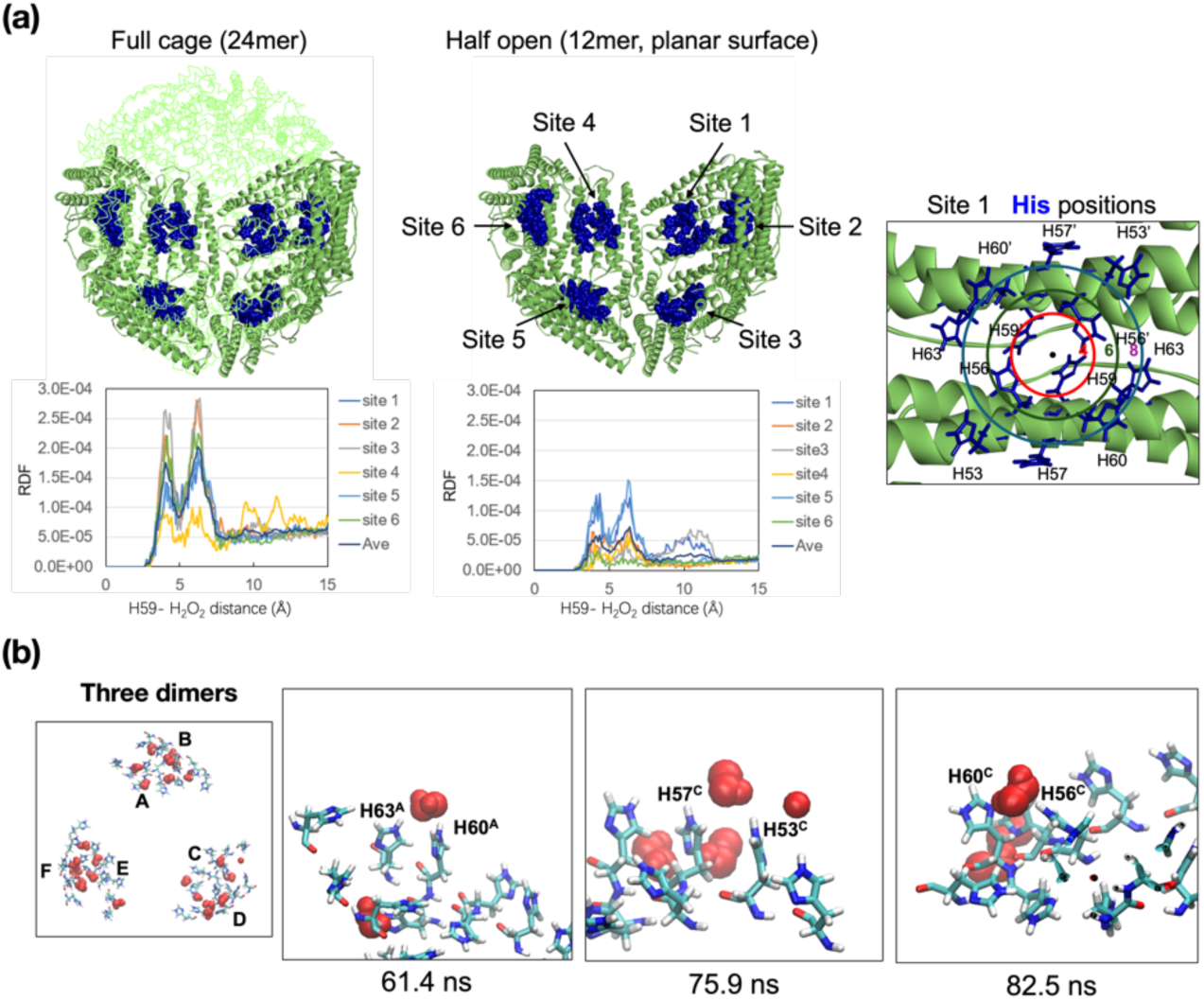
MD simulations of H_2_O_2_ in the **Fr-H6** cage and the confinement effect. (a) Binding of H_2_O_2_ to the His-clusters in 24-mer (full) and 12-mer (half) cages of **Fr-H6**. The full and half cages are shown at the top, and the close-up of the His-cluster is shown on the right, where the mutated His residues are represented by blue spheres and sticks, respectively, and the red, green and blue circles on the right represent the distances of 4, 6 and 8 Å from the center of mass of the H59 pair, respectively. The RDF values of H_2_O_2_ from the H59 pair are shown at the bottom. (b) Snapshot structures of H_2_O_2_ on the His-clusters in the simulation. The three adjacent His-clusters consisting of dimers AB, CD, EF are shown on the left, with H_2_O_2_ (red). Three snapshots are shown on the right, where H_2_O_2_ are located between His63^A^ and His60^A^ at 61.4 ns, between His57^C^ and His53^C^ at 75.9 ns, and between His60^C^ and His56^C^ at 82.5 ns, respectively.

Significant differences in the behavior of H_2_O_2_ within the **Fr-H8** cage compared to **Fr-H6** cages were observed. In **Fr-H6**, the RDF values of H_2_O_2_ at the 12 two-fold symmetric interfaces all have similar patterns (Figure 4a), whereas in **Fr-H8**, they were mainly divided into two major groups (Figure S12). The first pattern, observed at the six two-fold symmetric interfaces, has two peaks near 3 Å and 7 Å as in the Fr-H6 cage (Figure S12a). The second pattern, identified at four sites, showed a maximum near 7 Å but does not exhibit an apparent shoulder at 3 Å (Figure S12b). **Fr-H8**, a mutant of **Fr-H6** where R52 and R64 are additionally replaced with His, exhibits double conformations at His residues 53, 57, and 60, suggesting that the His residues in the His-cluster have greater flexibility compared to those in **Fr-H6** (Figure 2). Therefore, although the clusters in **Fr-H8** are larger than the clusters of **Fr-H6** and the frequency of H_2_O_2_ approaching the His-cluster increases, we postulate that **Fr-H8** is less likely to form specific sites binding with H_2_O_2_ compared to **Fr-H6**. Thus, the higher activity of the reaction between H_2_O_2_ and TMB in **Fr-H6** suggests that specific reactive regions are formed within the His-clusters in **Fr-H6**.

To analyze the approach of TMB molecules to the His-clusters, MD simulations of the **Fr-H6** system (containing 132 H_2_O_2_ molecules) with eight TMB molecules were performed (Figure S13). It was observed that TMB molecules remain close to the His-clusters for a longer time than H_2_O_2_ molecules (Figure S14). Among the TMB molecules numbered 1-8, TMB1 remains at site 1 of the 12 His-clusters for 16.9 ns, transitions to site 12 at 18.4 ns, and settles further into the region between the His dimers at site 12 after 38.7 ns (Figure S13). TMB3 initially remains at site 6 for 11.8 ns, transitions to site 12 for 9.5 ns (26.3-35.7 ns), but then returns to site 6 at 39.7 ns and remains at that site. Interestingly, TMB6 remains at site 3 for 30.5 ns, transitions to site 6 at 34.4 ns (until 65.7 ns) and then moves to site 7 at 78.9 ns and remains at that site until 94.4 ns. The TMB molecules appear to remain at the His-cluster sites for several tens of ns with repeated transitions.

An analysis of the approach of TMB and H_2_O_2_ molecules to the His-cluster indicates variations where the TMB molecule contacts the H_2_O_2_ molecule interacting with two His side chains (TMB 1 at 10.1 ns, TMB 3 at 58.6 ns, and TMB 6 at 79.4 ns, as shown in Figure S13 (close-ups in each figure). In the previous study, the oxidation reaction of TMB molecules which involves H_2_O_2_ interacting with His residues was demonstrated using theoretical analysis.^29^ Therefore, as observed above, it appears that the reaction proceeds more frequently in the **Fr-H6** system due to the multiple instances of approach of TMB molecules to the His-clusters.

## Discussion

This work demonstrates that an array of His residues on the B-helix of ferritin leads to the formation of His-clusters at the two-fold symmetric interface. These His residues interact with the corresponding His residues on the B-helix of the opposite subunit to form inter-subunit interactions. Thus, the His-cluster exhibits enzyme-mimetic oxidase activities by catalyzing one-electron oxidation of TMB in the presence of H_2_O_2_. In previous investigations, His residues were introduced within short peptides to assemble into supramolecular nanostructure, exhibiting similar enzyme-mimetic oxidase activities.^26,29,35^ In such cases, only the residues exposed on the outer surface of the structure must be maintained. In contrast, ferritin offers the unique advantage of the cage structure.^9^

Considering kinetic parameters, the catalytic activity of ferritin mutants follows the order **Fr-H6** > **Fr-H8** >> **Fr-H4in** > **Fr-H2** > **Fr-H4ex** (Table 2). **Fr-H6** which has two fewer His residues, has more than double the activity of **Fr-H8** (*k*_cat_/*K*_m_, 25.47 *vs.* 57.95). The X-ray structure of **Fr-H6** has two Arg residues at positions 52 and 64 which are replaced by His residues in **Fr-H8** (Figure. 2b). His52 and His64 form a long inter-subunit interaction through His63 in **Fr-H8**. We postulate that the positively charged Arg residues at positions 52 and 64 in **Fr-H6** might trigger the substrate interaction to generate the highest catalytic activity. This is correlated with the lower activity of **Fr-H4ex** and **Fr-H4in** where Arg52 and Arg64 are each replaced by a His residue (*k*_cat_/*K*_m_, 5.53 and 8.25). In an attempt to identify the minimal requirement for catalysis at the active site, it was found that the **Fr-H2** mutant, with just three His residues at positions 49, 53, and 57 (Figure. 2e), has activity similar to the activities of **Fr-Hex** and **Fr-Hin** (*k*_cat_/*K*_m_ of 7.22 vs 5.53 and 8.25 M^−1^s^−1^). This suggests that a set of three His residues, including a flexible middle His (His53 with double conformations), may be one of the smallest units required to promote peroxidase activity. However, other factors such as inter-subunit interactions, the number of His residues, and surrounding residues also appear to play significant roles.

To investigate the role of histidine in more detail, we estimated the overall activity (*k*_cat_/*K*_m_) per histidine residue present in the B-helix of ferritin mutants (Table S4). **Fr-H6** has the highest value of 0.35 M^−1^s^−1^ per histidine in the B-helix. The activity does not increase linearly with the number of histidine residues, suggesting that the position of histidine matters more than the number of histidine residues. Compared with an oligohistidine^26^ containing 15 histidine residues, the activity value of **Fr-H6** is 7.5-fold greater than that of oligohistidine for TMB. Although this is a rough estimation based on the number of histidine residues, it does not account for the nature and the effect of the assembly structure. For example, our MD simulation revealed that both TMB and H_2_O_2_ frequently contact the His cluster within the cage, and the frequency decreases when the cage volume is restricted to half. This indicates that the confined protein environment enhances the activity and is consistent with the previous reports.^36–38^ The small TMB molecule can easily enter the cage through the three-fold channels.^39^ In considering all of the present observations, it appears that the exposed His-cluster on the internal surface of the unique ferritin cage structure plays a significant role in promoting high catalytic activity. Although we achieved significant peroxidase activity in the present ferritin mutants, the overall activity (*k*_cat_/*K*_m_) of **Fr-H6** is much lower than that of a representative native peroxidase enzyme (57.95 *vs*. 9.22×10^6^ M^−1^s^−1^).^40^

### Conclusions

In summary, we successfully constructed His-clusters at the two-fold symmetric interface of ferritin and demonstrated oxidase-mimetic catalytic oxidation of TMB with H₂O₂. The high-resolution X-ray structures revealed His-His interactions (5–6Å) in the designed mutants. The Fr-H6 mutant efficiently catalyzes TMB oxidation via Michaelis-Menten kinetics with a higher affinity for TMB than for H₂O₂. MD simulations revealed that both substrates interact frequently with the His cluster, exhibiting stronger interactions within the confined cage as a result of the cage effect. Our findings emphasize that the number and positioning of His residues, as well as inter-subunit interactions significantly enhance the catalytic activity. These insights into engineered His-clusters are promising and are expected to be incorporated into bioinspired catalysts. This engineering strategy is expected to be useful in development of artificial metal-free enzymes for use in sustainable chemical production, biomaterials, and environmental remediation.

## Acknowledgements

This work was supported by JSPS KAKENHI (Grant Nos. 22H00347 to T.U., and 22H04744 to B.M.) from Ministry of Education, Culture, Sports, Science and Technology, the Adaptable and Seamless Technology Transfer Program through Target-driven R&D (JPMJTR224A) from the Japan Science and Technology Agency to T.U., and JST SPRING (JPMJSP2106 and JPMJSP2180 to J. T.).

## Contributions

J. T., B.M., T.P. and T.U. designed the research. J.T. carried out the wet experiments. T. F. conducted the MD simulation. All authors analyzed the data, discussed the results and co-wrote the paper.

