## Supplementary Information for "An artificial metal-free peroxidase designed using a ferritin cage"

Jiaxin Tian[1], Basudev Maity[1*], Tadaomi Furuta[1*], Tiezheng Pan[1], Takafumi Ueno[1,2*]

[1] School of Life Science and Technology, Institute of Science Tokyo, 4259 Nagatsuta-cho, Midori-ku, Yokohama, Kanagawa 226-8501, Japan
[2]Department of Life Science and Technology, Research Center for Autonomous Systems Materialogy (ASMat), Institute of Integrated Research, Institute of Science Tokyo, 4259 Nagatsuta-cho, Midori-ku, Yokohama, Kanagawa 226-8501, Japan

**Table of Contents**

**Experimental section**
**Figures S1-S14**
**Tables S1-S4**
**Description of Supplementary Movies S1-S2**
**References**

### Experimental section

### Materials and method

All the materials such as 3,3',5,5'-Tetramethylbenzidine (TMB), Hydrogen peroxide etc. were purchased from commercial sources like Wako, TCI etc. and used at it received. The absorption spectral measurements were performed using UV-2600PC UV-vis spectrometer (Shimazu).

### Protein expression and purification

Recombinant L-chain horse spleen apo-ferritin (rHLFr) was used for the study. Mutants were prepared by inverse polymerase chain reaction (iPCR) method and the primers are listed in Table 1. Protein expression was carried out in Novagen's NovaBlue competent cells, which were transformed with the pMK2 expression vector.[1] After expression, the protein was isolated by sonication, heating at 65°C for 15 min followed by purification through anion exchange chromatography (Q-Sepharose) and size exclusion chromatography (S-300 Sephacryl). The purity and assembly of the 24-mer ferritin cage were confirmed by native polyacrylamide gel electrophoresis (PAGE), and the molecular mass of the monomers was determined using matrix assisted laser desorption ionization-time of flight mass spectrometry (MALDI-TOF-MS) (Bruker ultrafle Xtreme). Protein concentrations was determined based on the molar absorption coefficient value of $4.6\times10^5$ $M^{-1}cm^{-1}$.

### Protein crystallization

The crystallization of ferritin and its mutants was performed through the hanging drop vapor diffusion technique.[2] Crystallization drops consisting of 1.5 μl of protein solution (20-30 mg/ml, 50 mM Tris-HCl, pH 8.0, 150 mM NaCl) mixed with 1.5 μl of precipitant solution containing 0.5–1.0 M $(NH_4)_2SO_4$ and 12.5–20.0 mM $CdSO_4$.The mixtures were incubated at 20°C. The crystals as long as 200-300 μm size appeared within one day.

### X-ray structure determination

The X-ray diffraction measurements were carried out using Rigaku XtaLaB Synergy-DW (Cu-Kα) diffractometer at Suzukakedai Materials Analysis Division, Tokyo Institute of Technology. Crystals were soaked in the cryoprotectant solution (1.0 M $(NH_4)_2SO_4$, 20 mM $CdSO_4$) containing 25% ethylene glycol for ~30 seconds before mounting for data collection. Data processing has been done automatically with CrysAlisPro software. Scaling was done using AIMLESS program in CCP4. The phase was determined using the MOLREP program in CCP4 using ferritin wild-type structure as a model (PDB: 1DAT).[3] The model was refined in REFMAC5 in CCP4.[4] The structures were rebuilt interactively in COOT followed by REFMAC5 refinement until a reasonable structure was obtained.[5] Due to insufficient electron density, the C-terminal residue Asp174 was left unassigned and the side chain of Lys172 was modeled as Ala in all structures. The positions of Cd ions were assigned based on the anomalous difference Fourier map at a cutoff of 4.0σ. The final model was validated through the wwPDB validation server and MolProbity analysis prior to deposition in the Protein Data Bank (PDB). Accession codes are given in Table S2. The structures were visualized using PyMOL (The PyMOL Molecular Graphics System, Version 2.1, Schrödinger, LLC).

### Circular dichroism (CD) spectroscopy

CD spectra of **Fr-H6** and **Fr-H8** were measured on the J-820 CD spectrometer (JASCO). A quartz cuvette with a path length of 0.1cm was used. The **Fr-H6** and **Fr-H8** samples with a concentration of 0.3 μM in 50mM Tris-HCl, pH 8.0, 150mM NaCl, were used for the study. The CD spectra were recorded in the far-UV region with a wavelength range of 190–260 nm at 25°C, averaging five scans with a bandwidth of 1 nm. CD temperature scans for **Fr-H6** and **Fr-H8** were performed at λ=222 nm, starting from 25 °C to 90 °C, with a thermal gradient of 2 °C /min.

### Peroxidase-like activity assays

To avoid any possibility of metal contamination, always deionized water was used, and ferritin samples were pretreated with EDTA before use. The purified ferritin was concentrated and incubated with 500 equiv. of EDTA for 2 hours with shaking at 500 rpm at 4 °C. Then dialyzed overnight at 4 °C against 2 L of 50 mM Tris-HCl buffer, pH 8.0, 150 mM NaCl. During the dialysis process, the buffer was changed twice to ensure the effective removal of unbound ions and impurities. Then, the ferritin samples were purified by size exclusion chromatography (S-300 Sephacryl column) using 50 mM Tris-HCl (pH8.0) containing 0.15 M NaCl as eluting buffer. For peroxidase-like activity, the reaction system contained 10 μM ferritin, 50 mM sodium acetate buffer (pH 4.0), 100-1000 μM of 3,3',5,5'-tetramethylbenzidine (TMB), and 2.5-20 mM $H_2O_2$ in a total volume of 0.2 mL. In a typical reaction, to the mixture of protein and TMB, $H_2O_2$ was added and subsequently, the absorbance at 652 nm was monitored with time in a UV–Visible spectrophotometer. Due to catalytic activity, the color of the solution was changed to blue. During measurement, the temperature was recorded as 29.5±1 °C.

### MD simulation

All the molecular dynamics (MD) simulations were conducted using the Amber 22 software package[6]. The structures of **Fr-H6** and **Fr-H8** were based on the crystal structures solved in this study [PDB 9KF9, 9KFA]. The Amber ff19SB parameters were used for proteins, and TIP3P model for water molecules. For TMB and hydrogen peroxide ($H_2O_2$), their partial ESP charges were first generated at the HF/6-31G* level using Gaussian 16 software[7], and then their force field parameters were generated through the antechamber module using RESP fitting and GAFF parameters. For the full-cage system with $H_2O_2$, 11 $H_2O_2$ molecules were first manually placed between the two histidines of the **Fr-H6** His cluster (11 $H_2O_2$ molecules per Fr dimer), and then replicated across all 12 His clusters (132 $H_2O_2$ molecules in total), and these same 132 molecules were also used in the **Fr-H8** system. For the half-cage system, these $H_2O_2$ molecules were kept, while the 6 Fr dimers (12 Fr molecules) were deleted. In the TMB-containing system, the two TMB molecules were first manually placed about 20 Å away from each H57 for one Fr dimer and then replicated to the full **Fr-H6** cage (12 Fr dimers), leaving 8 symmetric TMB molecules out of 24 (132 $H_2O_2$ and 8 TMB molecules in total). For each system, after solvation, 300 steps of energy minimization with heavy atoms restrained was performed, followed by 500 ps of NVT equilibration and 500 ps of NPT equilibration with the same restraints. Finally, a 100 ns production run was performed (where the main chain restraints were applied in place to account for the half-cage system, a quasi-planer system).

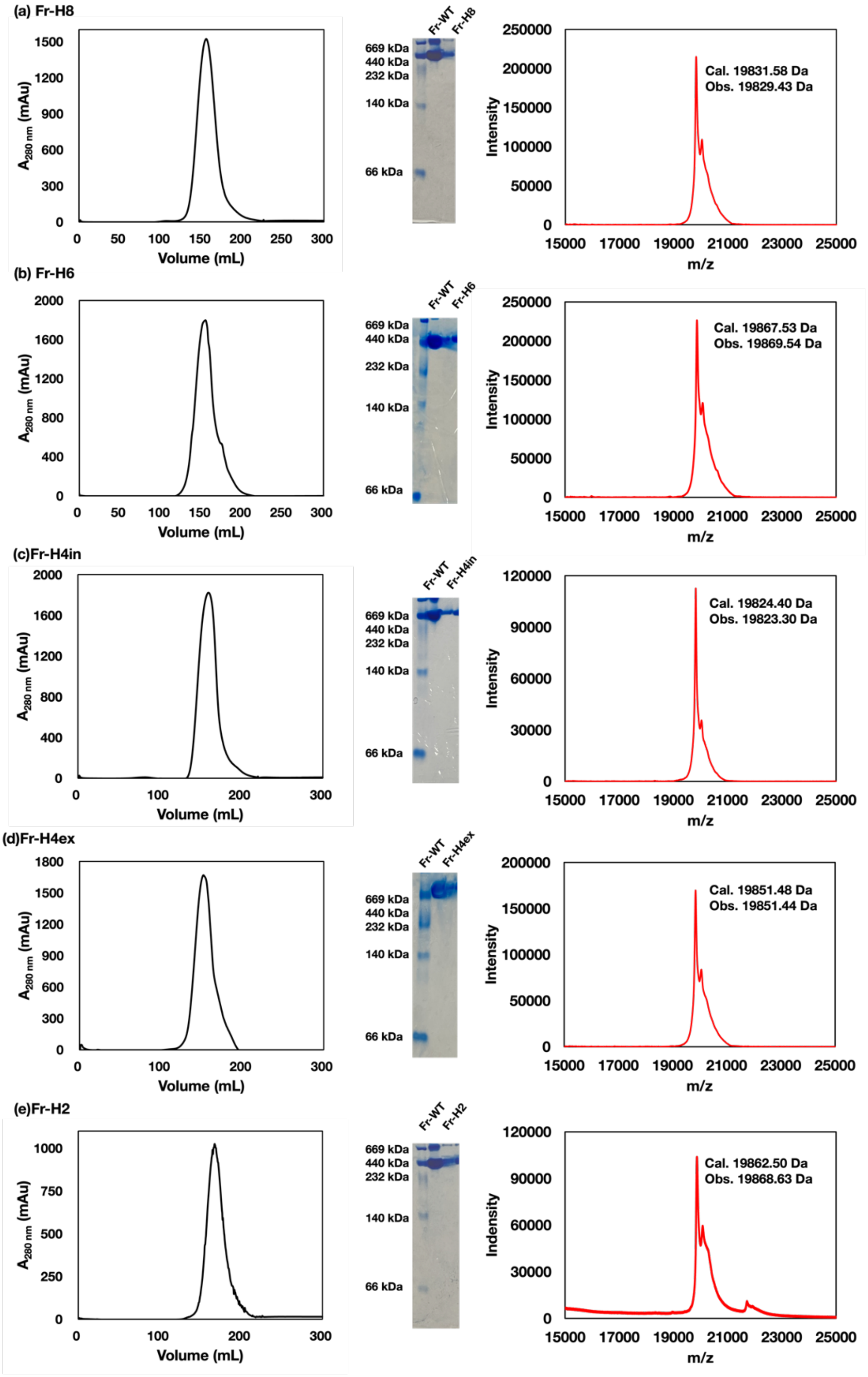


**Figure S1.** Characterization of ferritin mutants used in the study. SEC elution profiles , native PAGE images and MALDI-TOF-MS spectra of Fr mutants are shown for (a) **Fr-H8**, (b) **Fr-H6**, (c) **Fr-H4in**, (d) **Fr-Hex** and (e) **Fr-H2**. In the native PAGE images, the first lane represents molecular weight marker.

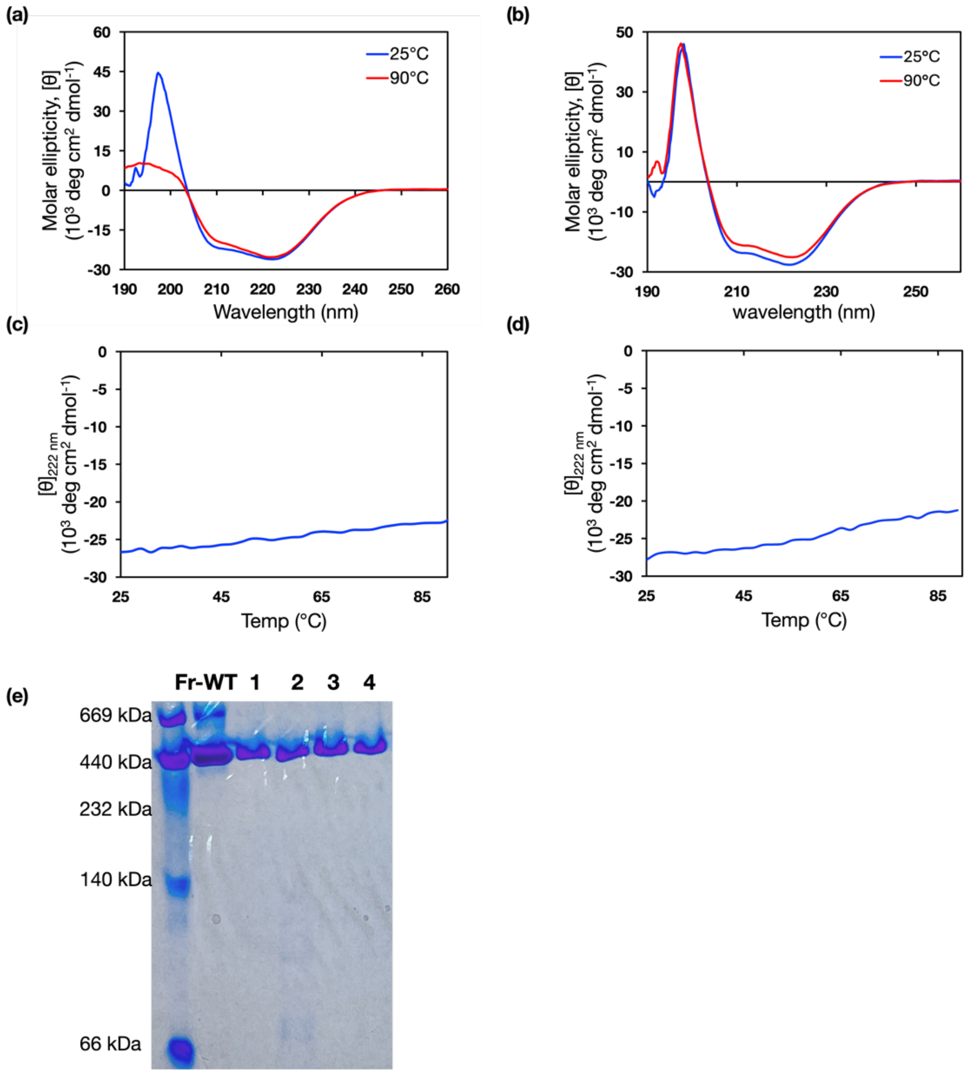


**Figure S2**. Stability of **Fr-H6** and **Fr-H8** by CD spectroscopy. CD spectra of (a) **Fr-H6** and (b) **Fr-H8** measured at 25 °C (before heating) and 90 °C (after heating) in 50 mM Tris-HCl, 150 mM NaCl, pH 8.0. Changes of molar ellipticity at 222 nm, $[\theta]_{222}$, of (c) **Fr-H6** and (d) **Fr-H8** during heating from 25 °C to 90 °C with a temperature gradient of 2 °C $min^{-1}$. (e) Native PAGE image of **Fr-H6** and **Fr-H8** at pH 8.0. Lanes 1 and 2: **Fr-H6** and **Fr-H8** at 25 °C. Lanes 3 and 4: **Fr-H6** and **Fr-H8** after heating to 100 °C and then cooling to 25 °C.

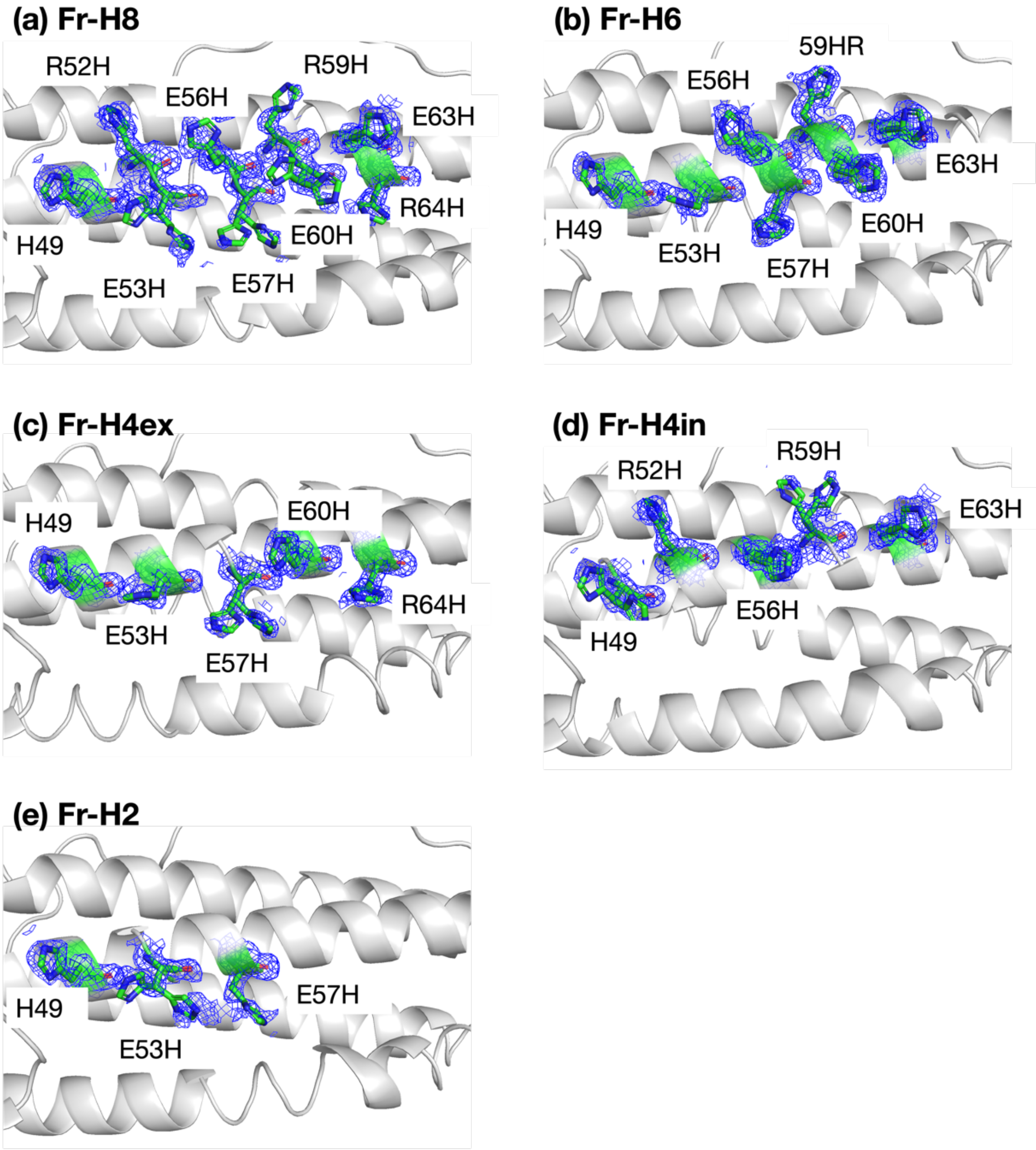


**Figure S3**. Electron density of the crystal structures of ferritin His mutants (a-e). $2F_o$-$F_c$ maps at 1σ are shown in blue mesh for His49 and other mutated histidine residues. The images presented are the screenshots from PyMol.

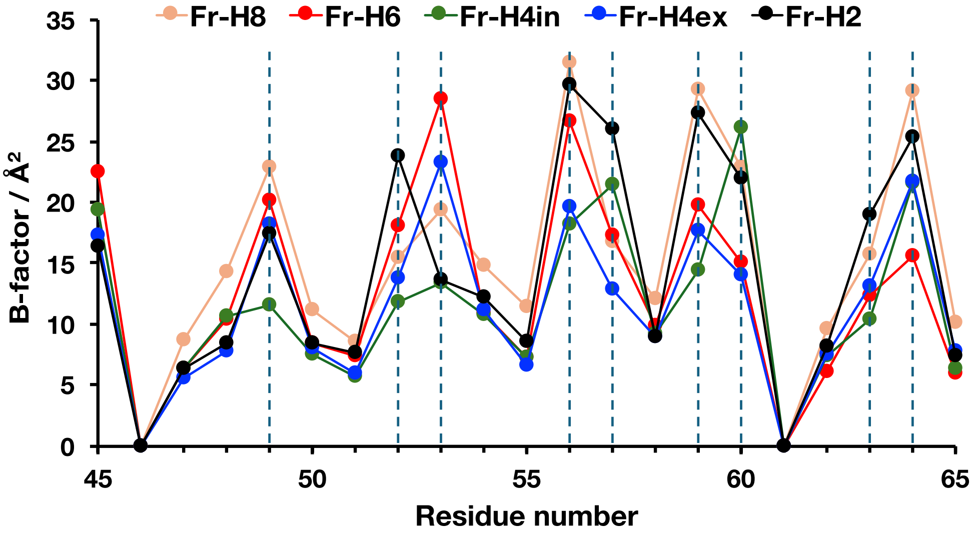


**Figure S4**. Sidechain B-factors (Å$^2$) of ferritin mutants for residues 45-65. The dashed vertical lines represent the residue positions at which His mutations were done in the respective ferritin mutants. The zero B-factor represents Gly residue with no sidechain.

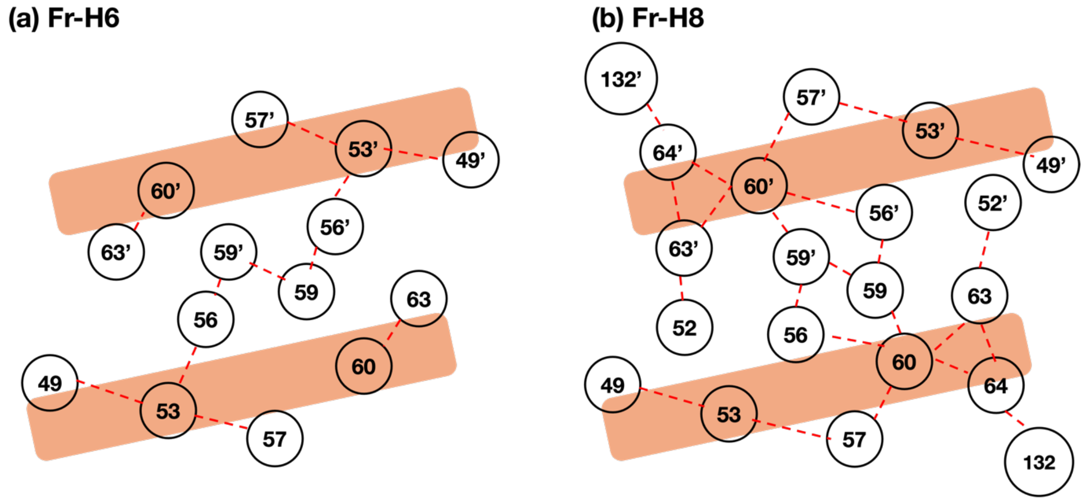


**Figure S5.** Schematic image of His-cluster in **Fr-H6** (a) and **Fr-H8** (b). The histidines with double conformations are also shown using a single symbol

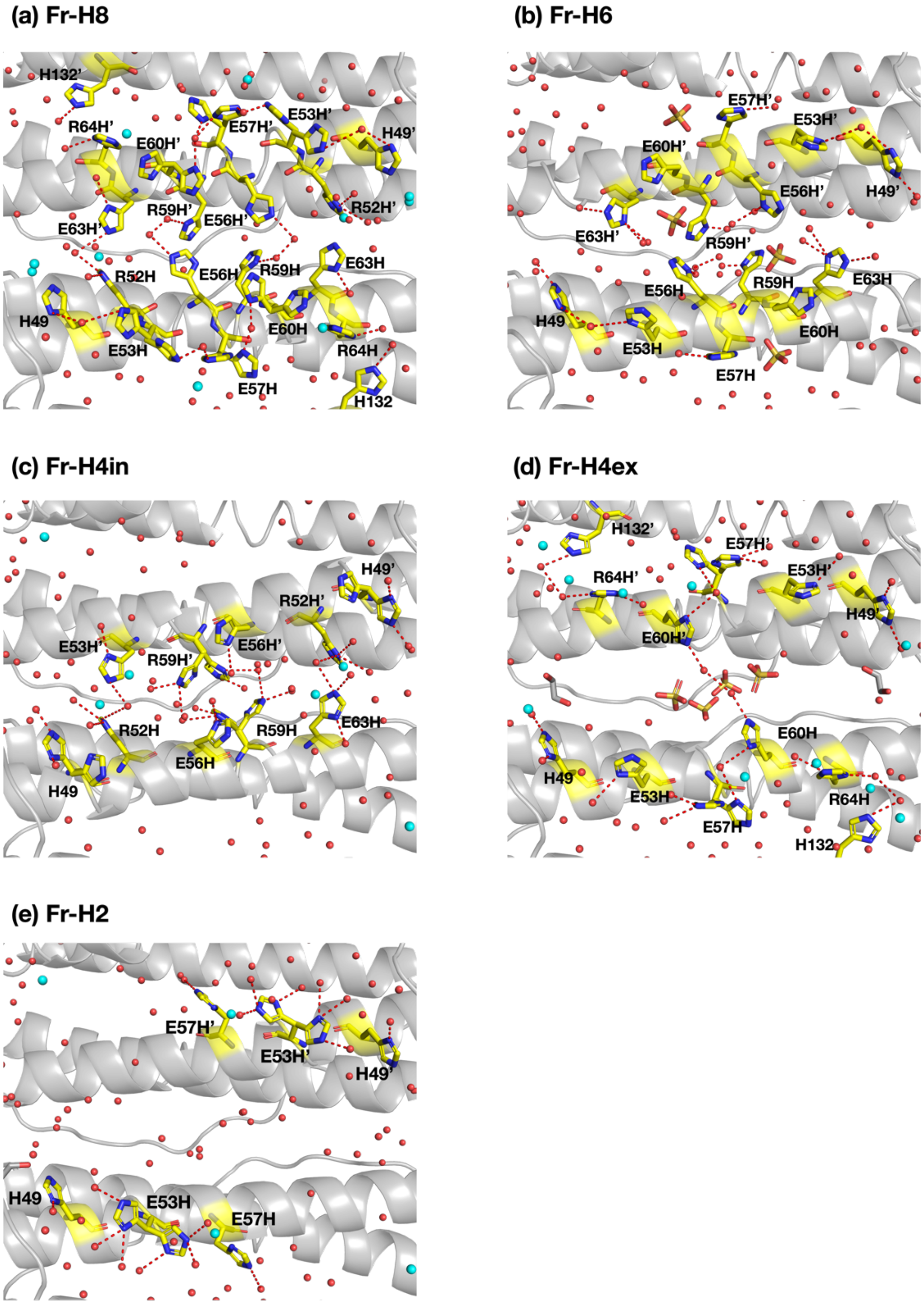


**Figure S6**. Water molecule networks surrounding the histidine clusters in various ferritin mutants. Water molecules are shown in red sphere. The Cd (cyan sphere) and sulfate (orange and red stick) ions come from the precipitant $CdSO_4$ and $(NH_4)_2SO_4$ used for crystallization. The ethylene glycol (gray and red stick) comes from the cryoprotectant.

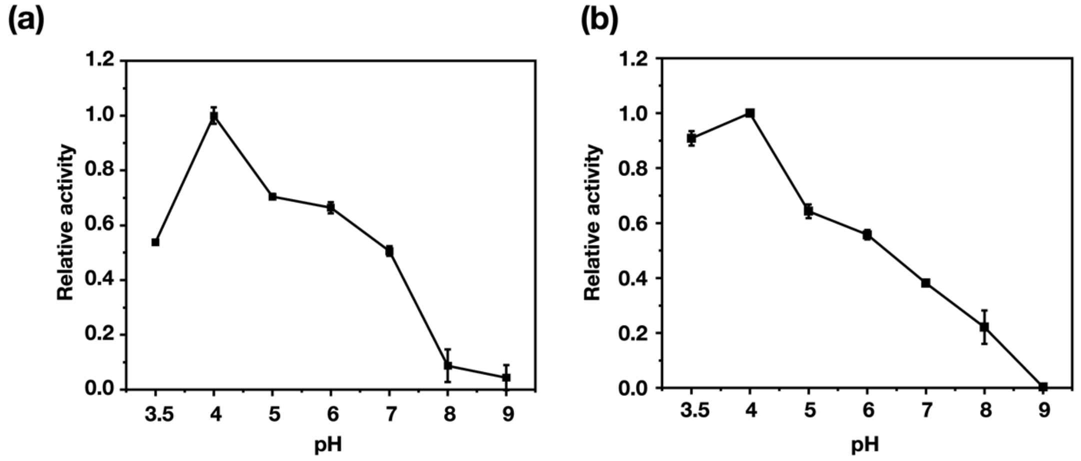


**Figure S7**. Effect of pH on the relative activity of (a) **Fr-H8** and (b) **Fr-H6** mutant. The relative activity was calculated based on the change in absorbance at 652 nm over 3 minutes using TMB as the substrate, reflecting the reaction velocity of the peroxidase-like activity under different pH conditions. Reaction condition: [protein] = 10 µM, [TMB] = 700 µM, [$H_2O_2$] = 20 mM. Buffer: 50 mM NaOAc (pH 3.5-6.0); 50 mM MES, 0.5 M NaCl (pH 7.0); 50 mM Tris-HCl, 0.15M NaCl (pH 8.0-9.0).

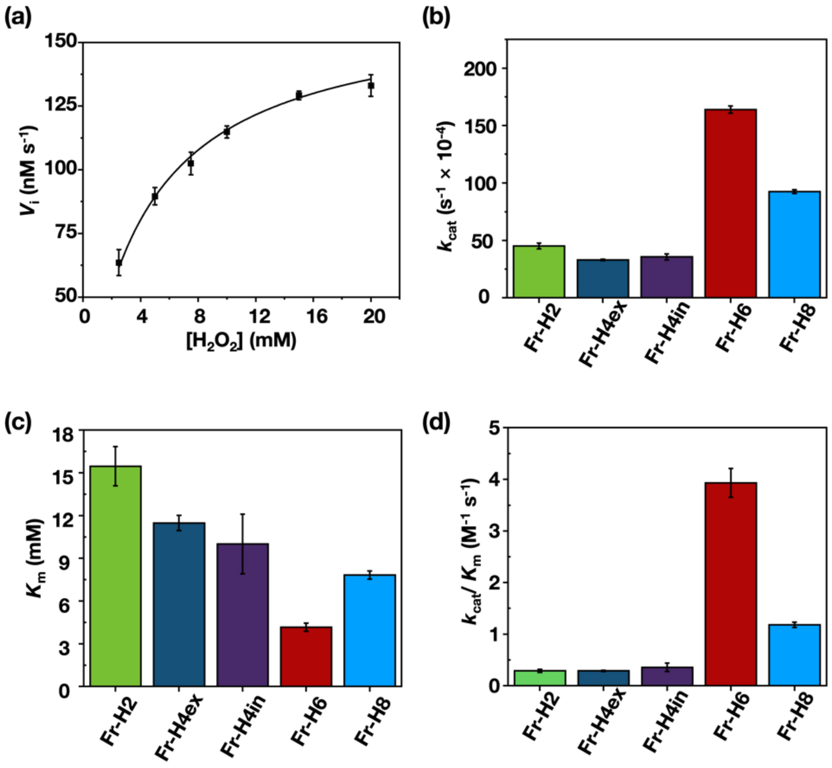


**Figure S8.** Evaluation of oxidase mimetic activities by Fr-His mutants. (a) Changes of reaction velocity ($V_i$) with increasing $H_2O_2$ concentration for the oxidation by **Fr-H6**. Reaction condition: [protein] = 10 μM, [TMB] =700 μM. (b) $k_{cat}$ (s$^{-1}$ ×10$^{-4}$), (c) $K_m$ (mM) and (d) $k_{cat}/K_m$ (M$^{-1}$ s$^{-1}$) for Fr-His mutants determined from the Michaelis–Menten curve. [protein] = 10 μM, [TMB] = 700 μM, in 50 mM NaOAc (pH 4.0).

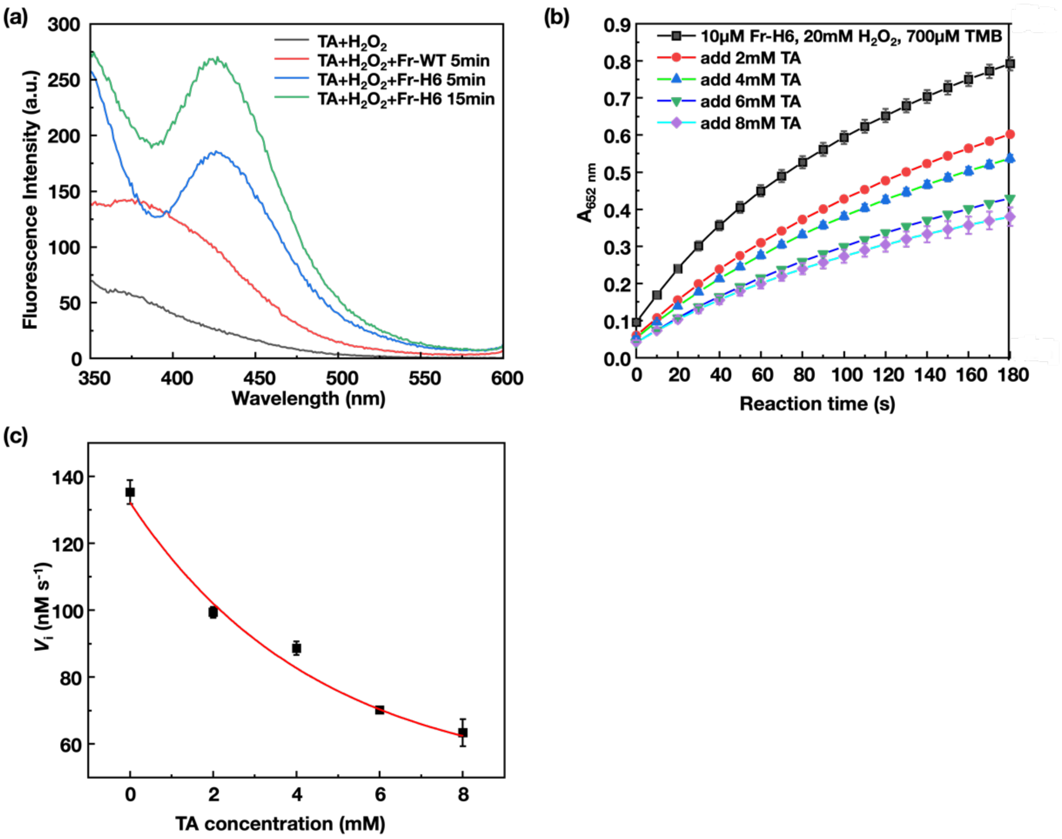


**Figure S9**. Effect of terephthalic acid (TA) on the peroxidase activity of **Fr-H6**. (a) Fluorescence spectra of terephthalic acid for **Fr-H6** after 5 and 15 min in the presence of $H_2O_2$, compared with FrWT. Reaction condition: [protein] = 10 μM, [$H_2O_2$] = 20 mM, [TA]= 0.25 mM. (b) Time-dependent absorbance changes at 652 nm for **Fr-H6** during the reaction in presence of various amount of TA. (c) Changes of reaction velocity ($V_i$) with increasing TA concentration for the oxidation by **Fr-H6**. [protein] = 10 μM, [TMB] = 700 μM, [$H_2O_2$] = 20 mM, in 50 mM NaOAc (pH 4.0).

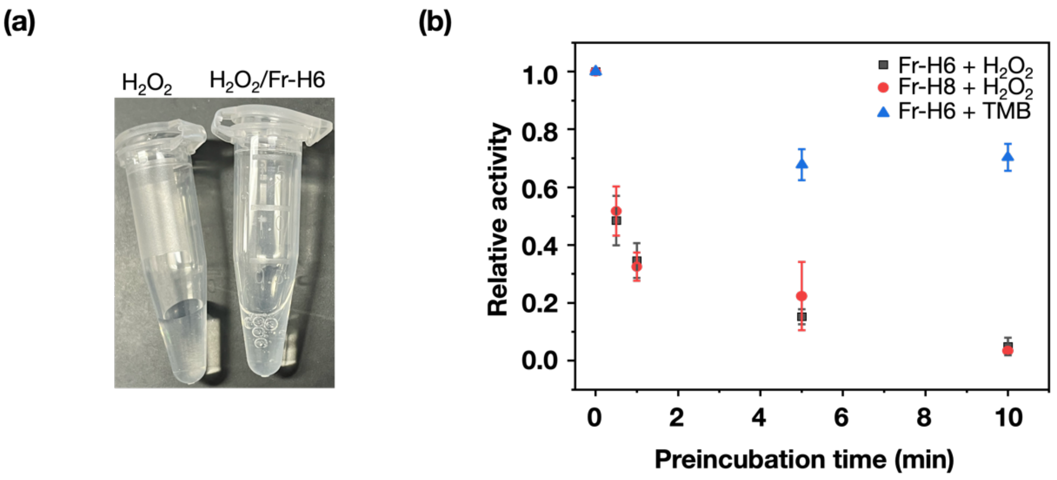


**Figure S10.** Catalase activity test with $H_2O_2$. (a) Optical images showing the bubble formation in the presence or absence of **Fr-H6** in $H_2O_2$ solution. Reaction condition: [protein] = 10 µM, [$H_2O_2$] = 2 M, in 50 mM NaOAc (pH 4.0). (b) Relative activity of **Fr-H6** and **Fr-H8** at different preincubation time with TMB or $H_2O_2$ under the same conditions as in (a). The relative activity was calculated based on the change in absorbance at 652 nm over 3 minutes using TMB as the substrate, reflecting the reaction velocity of the peroxidase-like activity. Blue triangle: Addition of $H_2O_2$ into the mixture of **Fr-H6** and TMB. Red circle: Addition of TMB into the mixture of **Fr-H8** and $H_2O_2$. Black square: Addition of TMB into the mixture of **Fr-H6** and $H_2O_2$.

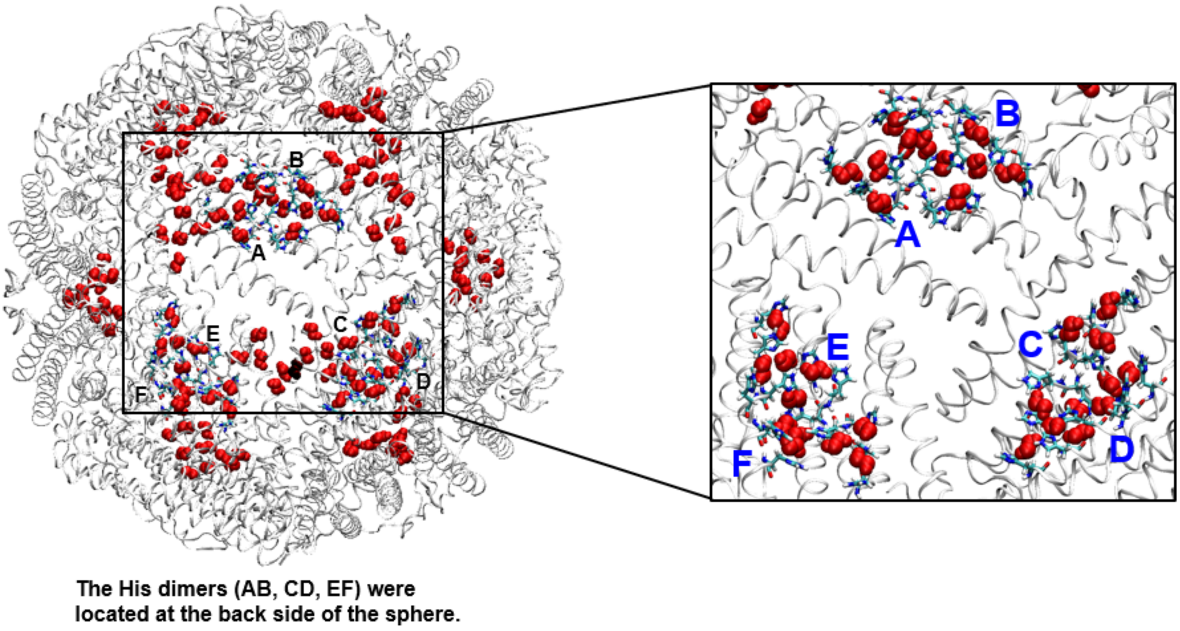


**Figure S11**: MD simulations of 24-mer **Fr-H6** with 132 $H_2O_2$ (initial structure). The dynamic behaviors of $H_2O_2$ can be found in Movie S2.

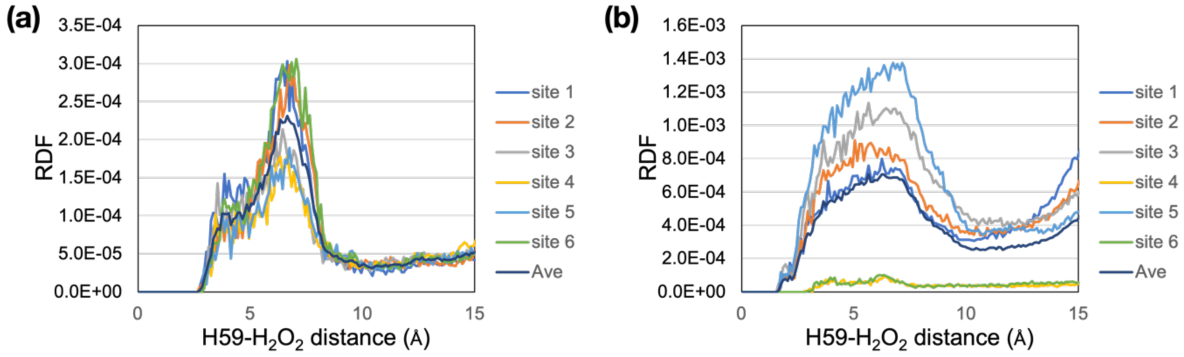


**Figure S12**: The radial distribution functions (RDFs) of $H_2O_2$ from H59 pair in **Fr-H8** simulation. (a) First pattern of RDF with two peaks around 3 Å and 7 Å. (b) Second pattern of that with a maximum of around 7 Å (without an apparent shoulder at 3 Å).

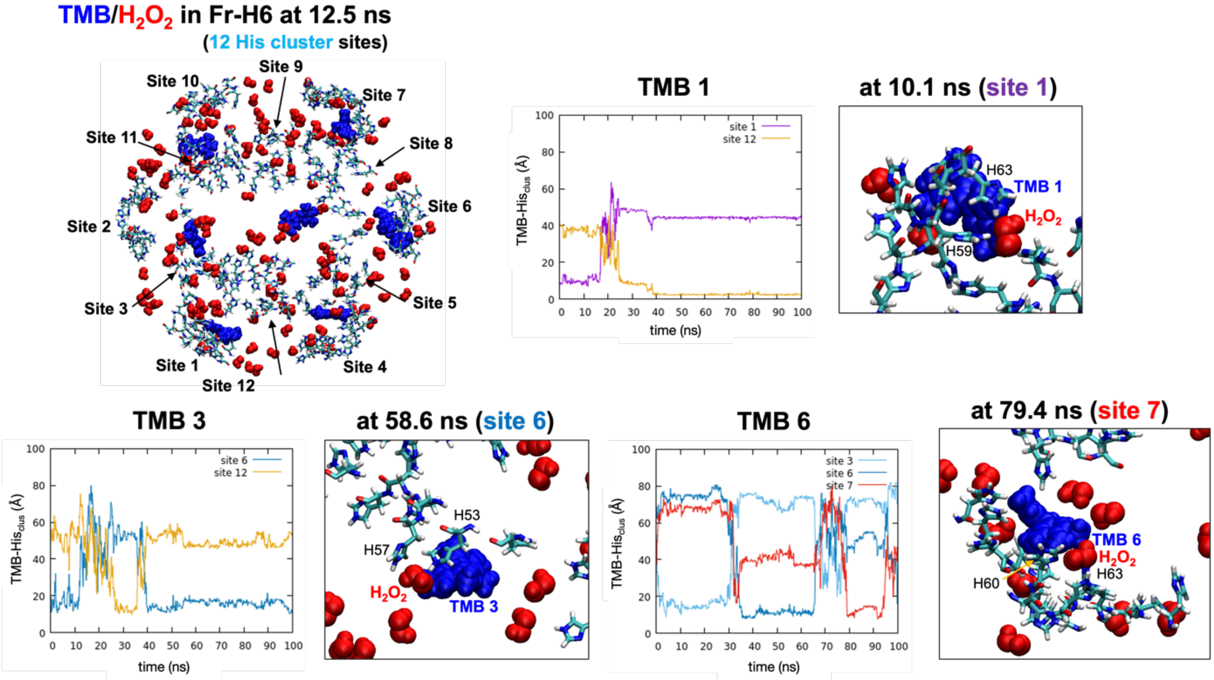


**Figure S13**: Molecular dynamic simulation of **Fr-H6** with TMB and $H_2O_2$. The entire cage (**Fr-H6**) and the His-cluster sites are shown in the upper left, where TMB and $H_2O_2$ molecules are represented by blue and red spheres. For the molecules TMB1, TMB3, and TMB6, the distances between the TMB molecule and the His-cluster and the snapshot structures are shown, respectively, where the TMB-His-cluster distance was calculated as the distance between the centers of mass of the TMB molecule (central two carbons) and H59 ($C_\beta$) pair of the ferritin dimer.

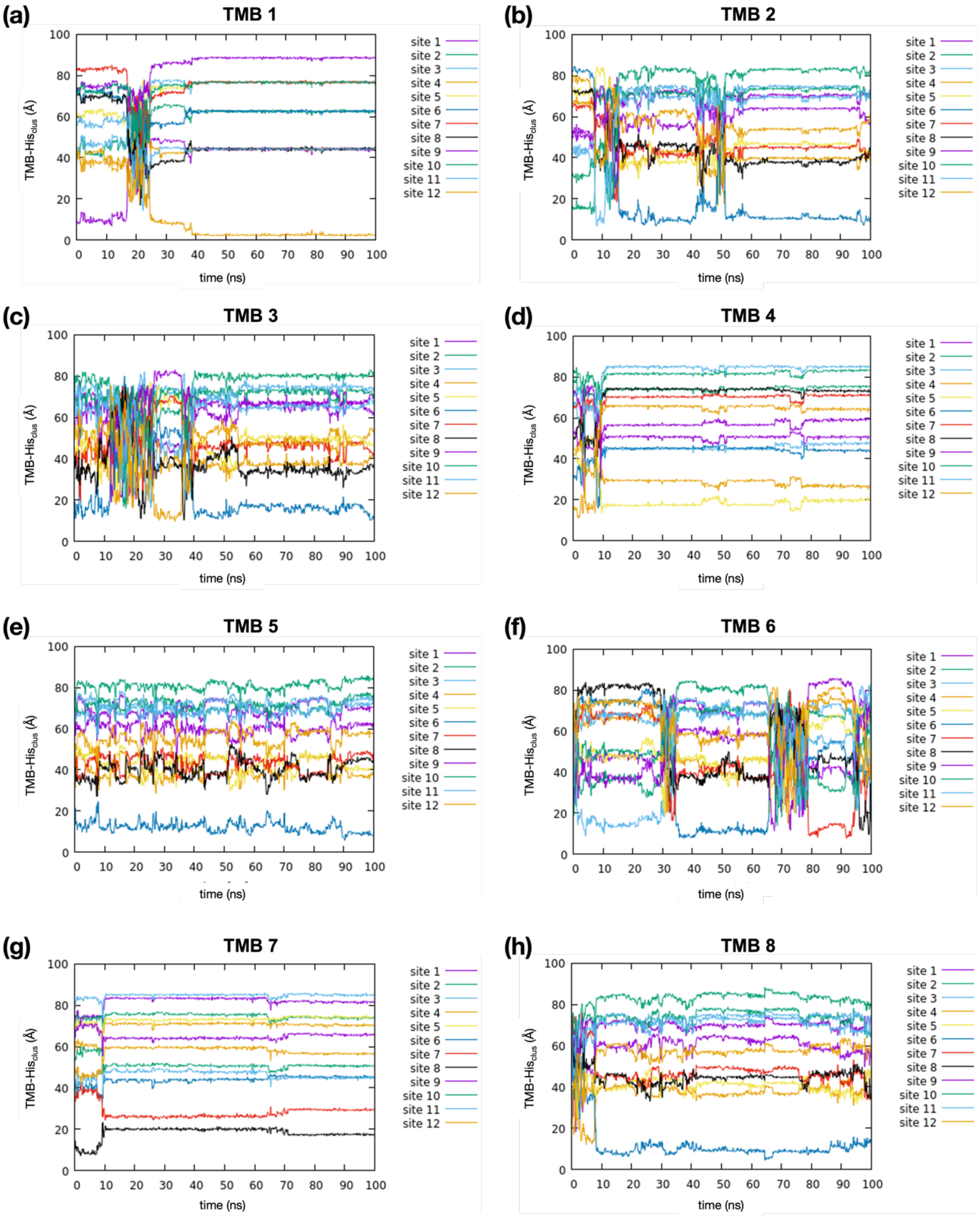


**Figure S14**: Changes in the distances between TMB and the His-clusters (H59 pairs) in **Fr-H6** simulation.

**Table S1A**: Fr-WT gene sequence used for expression in *E. coli*.

| |
|---|
| ATG AGC TCC CAG ATT CGT CAG AAT TAT TCT ACT GAA GTG GAG GCC GCC GTC AAC CGC CTG GTC AAC CTG TAC CTG CGG GCC TCC TAC ACC TAC CTC TCT CTG GGC TTC TAT TTC GAC CGC GAC GAT GTG GCT CTG GAG GGC GTA TGC CAC TTC TTC CGC GAG TTG GCG GAG GAG AAG CGC GAG GGT GCC GAG CGT CTC TTG AAG ATG CAA AAC CAG CGC GGC GGC CGC GCC CTC TTC CAG GAC TTG CAG AAG CCG TCC CAG GAT GAA TGG GGT ACA ACC CTG GAT GCC ATG AAA GCC GCC ATT GTC CTG GAG AAG AGC CTG AAC CAG GCC CTT TTG GAT CTG CAT GCC CTG GGT TCT GCC CAG GCA GAC CCC CAT CTC TGT GAC TTC TTG GAG AGC CAC TTC CTA GAC GAG GAG GTG AAA CTC ATC AAG AAG ATG GGC GAC CAT CTG ACC AAC ATC CAG AGG CTC GTT GGC TCC CAA GCT GGG CTG GGC GAG TAT CTC TTT GAA AGG CTC ACT CTC AAG CAC GAC TAA |

**Table S1B**: Sequence of primers used to generate Histidine cluster.

| |
|---|
| **Fr-H2**<br>5’-CGCCACTTGGCGGAGCACAAGCGCGAGGGTGC -3’(Forward)<br>5’- GAAGAAGTGGCATACGCCCTCCAGAGCC-3’(Reverse) |
| **Fr-H4in**<br>5’-CACGAGTTGGCGCACGAGAAGCACGAGGGTGCCCACCGTCTCTTGAAG -3’(Forward)<br>5’- GAAGAAGTGGCATACGCCCTCCAGAGCCACATCGTC-3’(Reverse) |
| **Fr-Hex**<br>5’- CGCCACTTGGCGGAGCACAAGCGCCACGGTGCCGAGCATCTCTTGAAGATGC-3’(Forward)<br>5’- GAAGAAGTGGCATACGCCCTCCAGAGCCACATCGTC-3’(Reverse) |
| **Fr-H6**<br>5’-CGCCACTTGGCGCACCACAAGCACCACGGTGCCCACCGTCTCTTGAAGATGCAAAACCAGCGC-3’(Forward)<br>5’- GAAGAAGTGGCATACGCCCTCCAGAGCCACATCGTC-3’(Reverse) |
| **Fr-H8**<br>5’-CACCACTTGGCGCACCACAAGCACCACGGTGCCCACCATCTCTTGAAGATGCAAAACCAGCGC -3’(Forward)<br>5’- GAAGAAGTGGCATACGCCCTCCAGAGCCACATCGTC-3’(Reverse) |

**Table S2**: Selected crystallographic parameters and refinement statics.

| **Dataset** | **Fr-H2** | **Fr-H4in** | **Fr-Hex** | **Fr-H6** | **Fr-H8** |
|---|---|---|---|---|---|
| PDB code | 9KF5 | 9KF7 | 9KF8 | 9KF9 | 9KFA |
| **Data collection** | | | | | |
| X-ray source | Cu Kα2 | Cu Kα2 | Cu Kα2 | Cu Kα2 | Cu Kα2 |
| Wavelength (Å) | 1.543 | 1.543 | 1.543 | 1.543 | 1.543 |
| Space group | F432 | F432 | F432 | F432 | F432 |
| Cell dimensions | | | | | |
| a = b = c (Å) | 182.68 | 181.70 | 180.93 | 181.64 | 182.00 |
| $\alpha = \beta = \nu$ (°) | 90.00 | 90 | 90 | 90 | 90 |
| Resolution limit (Å) | 23.81-1.50 (1.66-1.63) | 23.66-1.50 (1.53-1.50) | 20.23-1.5 (1.53-1.50) | 22.19-1.50 (1.53-1.50) | 22.23-1.57 (1.60-1.57) |
| Unique reflections | 32905 (1599) | 41552 (2012) | 41042 (1994) | 41233 (2016) | 36343 (1744) |
| Multiplicity | 7.8(5.6) | 7.1(4.7) | 8.7(5.7) | 7.2(4.9) | 9.4(6.8) |
| Completeness (%) | 99.4(100.0) | 100.0(99.6) | 99.9(99.8) | 99.3(99.6) | 99.5(100.0) |
| Mean (I /sigma(I)) | 16.1(2.1) | 18.1(2.1) | 21.4(2.8) | 17.0(2.2) | 17.3(2.1) |
| $R_{meas}$ | 0.110(0.769) | 0.095(0.759) | 0.076(0.671) | 0.092(0.698) | 0.092(0.717) |
| $R_{merge}$ | 0.102(0.697) | 0.088(0.673) | 0.072(0.608) | 0.085(0.623) | 0.087(0.662) |
| $R_{pim}$ | 0.039(0.322) | 0.035(0.345) | 0.025(0.276) | 0.033(0.310) | 0.029(0.271) |
| Half set correlation CC(1/2) | 0.996(0.796) | 0.998(0.760) | 0.999(0.819) | 0.996(0.804) | 0.997(0.861) |
| Average mosaicity (°) | 0.62 | 0.72 | 0.69 | 0.74 | 0.75 |
| Wilson B factor (Å$^2$) | 9.074 | 6.733 | 6.648 | 7.205 | 10.219 |
| **Refinement** | | | | | |
| Resolution (Å) | 1.63 | 1.50 | 1.50 | 1.50 | 1.57 |
| No. reflections used | 39521 | 39387 | 38967 | 38758 | 39295 |
| R-factor/R-free | 0.2174/ 0.2384 | 0.1665/ 0.1851 | 0.1564/ 0.1706 | 0.2062/ 0.2250 | 0.2149/ 0.2422 |
| **B-factors (Å$^2$)** | | | | | |
| Overall (protein part) | 12.071 | 10.562 | 10.906 | 10.392 | 13.547 |
| Main chain | 10.183 | 8.868 | 9.020 | 8.787 | 11.517 |
| Side chains | 13.755 | 12.074 | 12.545 | 11.820 | 15.305 |
| **R.m.s. deviations** | | | | | |
| Bond lengths (Å) | 0.0105 | 00126 | 0.0133 | 0.0122 | 0.0105 |
| Bond angles (°) | 1.8152 | 1.9133 | 1.9567 | 1.9281 | 1.8258 |
| **Ramachandran plot statics (%)** | | | | | |
| Favored region | 97.5 | 97.5 | 97.4 | 97.0 | 97.0 |
| Allowed region | 2.5 | 2.5 | 2.6 | 3.0 | 3.0 |
| Outlier | 0 | 0 | 0 | 0 | 0 |

Note: Values in the parentheses are for the highest-resolution shell. $R = \Sigma||F_o|-|F_c||/\Sigma|F_o|$, where $F_o$ and $F_c$ are the observed and calculated structure factor amplitudes, respectively. $R_{free}$ : R factor calculated on a partial set that is not used in the refinement of the structure. Ramachandran plot parameters were calculated using RAMPAGE.

**Table S3. The His-cluster in ferritin histidine mutants and the distance (Å) of His-cluster.**

| His-cluster | **Fr-H2** | **Fr-H4ex** | **Fr-H4in** | **Fr-H6** | **Fr-H8** |
|---|---|---|---|---|---|
| H49-53H | 6.2 | 6.7 | - | 6.7 | 6.1 |
| 52H-63H’ | - | - | 5.4 | - | 5.5 |
| 53H-56H | - | - | - | 6.5 | - |
| 53H-57H | 5.6 | 7.2 | - | 7.4 | 4.4 |
| 56H-59H’ | - | - | - | 7.0 | 6.6 |
| 56H-60H | - | - | - | - | 6.4 |
| 57H-60H | - | 6.6 | - | - | 6.3, 6.6 |
| 59H-59H’ | - | - | 5.1 | 5.8 | 6.1 |
| 59H-60H | - | - | - | - | 7.0 |
| 60H-63H | - | - | - | 5.2 | 6.1 |
| 60H-64H | - | - | - | - | 4.5 |
| 64H-H132 | - | 5.2 | - | - | 4.7 |
| 63H-64H | - | - | - | - | 6.8 |

‘-’: means that the histidine in this mutant is not contained in the His-cluster.
**His-cluster** (as defined here): The distance between the centers of the imidazole rings of two histidines is between 4 Å and 7.5 Å (and they are linked/clustered).

**Table S4. Comparison of the catalytic performance per histidine in the ferritin mutants.**

| Catalyst | No. of histidine per cage[a] | Substrate | $k_{cat}/K_m$ /histidine ($M^{-1}$ $s^{-1}$)[c] |
|---|---|---|---|
| FrWT | 24 | TMB | ND |
| **Fr-H2** | 72 | TMB | 0.1002 |
| **Fr-H4ex** | 144[b] | TMB | 0.0384 |
| **Fr-H4in** | 120 | TMB | 0.0688 |
| **Fr-H6** | 168 | TMB | 0.3450 |
| **Fr-H8** | 240[b] | TMB | 0.1061 |
| oligohistidine | 15 | TMB | 0.0467 |
| FrWT | 24 | $H_2O_2$ | ND |
| **Fr-H2** | 72 | $H_2O_2$ | 0.0040 |
| **Fr-H4ex** | 144[b] | $H_2O_2$ | 0.0020 |
| **Fr-H4in** | 120 | $H_2O_2$ | 0.0030 |
| **Fr-H6** | 168 | $H_2O_2$ | 0.0234 |
| **Fr-H8** | 240[b] | $H_2O_2$ | 0.0049 |
| oligohistidine[d] | 15 | $H_2O_2$ | 0.0020 |

Note: [a]FrWT monomer has six histidines (144 His per cage) and is catalytically inactive. Since all the mutations were made in the B-helix, only the histidines present in the B-helix were counted. FrWT has only one histidine (His49) in the B-helix (24 His per cage).

[b]Since H132 interacted with 64H in Fr-H4ex and Fr-H8, this residue was taken into account in the calculation.

[c]The 24-mer ferritin cage concentration was used for all the studies (Table 2 in the main text). The $k_{cat}/K_m$ values were divided by the number of histidines to obtain the value $k_{cat}/K_m$ / histidine.

[d]From reference 26 in the main text.

**Description of Supplementary Movies**

**Movie S1.** $H_2O_2$ behaviors within the 24-mer **Fr-H6** cage in the MD simulation. $H_2O_2$, His-clusters, and **Fr-H6** cage are represented in red sphere, stick, and ribbon model, respectively.

**Movie S2.** $H_2O_2$ behaviors at the 3-fold symmetric center in the MD simulation. $H_2O_2$ molecules within 4 Å of the His-clusters are visualized, and others are shown as in Movie **S1.**
